# Meningeal γδ T cells facilitate bacterial entry into the brain in neonatal meningitis and trigger long-term behavioural sequelae

**DOI:** 10.1101/2025.11.07.686929

**Authors:** Inês Lorga, Joana Soares, Joana Bravo, Pedro Mesquita, Nuno Ribeiro, João Rodrigues, Ahmed Hassan, Morgana Mesquita, Bruno Cavadas, Hristo Georgiev, Ana Magalhães, Teresa Summavielle, Manuel Vilanova, Julie C. Ribot, Elva Bonifácio Andrade

## Abstract

Neonatal bacterial meningitis is a life-threatening condition and a leading cause of neurodevelopmental impairment among survivors. Despite its prevalence, the role of meningeal immunity on disease pathology during early life remains largely unexplored. Using a clinically relevant mouse model of neonatal group B streptococcal meningitis and single-cell RNA sequencing, we observed that IL-17A (IL-17)-producing γδ T cells (γδ17 T cells) accumulate in the meninges during the acute phase of infection and persist throughout the lifespan. Importantly, mice deficient in γδ T cells or in IL-17 show a significantly lower bacterial colonisation in the brain parenchyma, and IL-17 neutralisation in the cerebrospinal fluid leads to a similar phenotype. Reduced blood-brain barrier permeability in the absence of γδ T cells results in decreased bacterial invasion and diminished microglia activation. Wild-type but not γδ T cell-deficient mice surviving infection exhibit increased hyperactivity and open space anxiety during early adulthood, a behavioural profile that may reflect attention deficit and hyperactivity (ADHD)-like tendencies. Altogether, these findings establish a pathogenic role for meningeal γδ17 T cells in early life, uncovering a key mechanism that drives long-term sequelae in GBS neonatal meningitis.

## Introduction

Group B *Streptococcus* (GBS), a commensal microorganism of the genitourinary and gastrointestinal tracts of adult humans, remains the leading agent causing bacterial meningitis in newborns^1^. GBS colonises the vagina or rectum of 15-40% of pregnant women^1^. Maternal vaginal colonisation is the primary risk factor for neonatal group b streptococcal infection, which can progress to meningitis^1^. Globally, this disease affects an infant every 3 minutes and kills one baby every 6-9 minutes^2,3^. GBS neonatal meningitis has an estimated mortality rate of 10-15% and results in life-long neurodevelopment impairment (NDI) in up to 50% of survivors^1^. Particular concern has arisen with some countries reporting a rising incidence of neonatal meningitis over the last decade^4–7^. The antibiotic prophylaxis administered to pregnant women vaginally colonised with GBS to prevent vertical transmission to the foetus is inefficient against meningitis ^8^, and no effective neuroprotective therapies are available to mitigate the long-term neurological sequelae.

The pathophysiology of GBS-induced neonatal meningitis and the mechanisms underlying its long-term NDI remain poorly understood despite significant efforts made in the past decades. Studies in animal models, such as haematogenous^9–11^ and direct brain GBS infection^12–15^, have faced challenges by generating data that inadequately replicate clinical phenotypes. This hindered the development of novel therapeutic interventions aiming at improving brain injury outcomes. To address these limitations, our group developed a mouse model of mother-to-progeny GBS transmission, which mirrors human infection pathogenesis^16^. Interestingly, this model revealed no leucocyte infiltration into the brain parenchyma or increased levels of inflammatory cytokines in the developing brain^16^. This challenges previous assumptions that the recruitment of immune cells into the parenchyma directly causes brain damage and NDI in GBS meningitis^17–20^.

In the past 20 years, the meninges have emerged as an immunologically active site rather than merely a physical protective barrier. The meninges harbour a wide repertoire of immune cells which play a role in haemostasis, infection and healing^21^. Additionally, meningeal immunity has been increasingly emphasised as crucial for maintaining steady-state brain function, particularly through the T cell compartment^22–30^. Cognitive functions, such as spatial learning and memory, and long-term potentiation depend on meningeal CD4^+^ and γδ T cells^22–27^. These cells are also important in regulating emotional behaviour, compulsive behaviour, nest building, anxiety and sociability^27–30^. Additionally, emerging evidence suggests that the meninges play essential roles in brain development, such as regulating the migration and positioning of neurons^31^. Despite the importance of the meninges, studies of neonatal meningitis have largely overlooked this anatomical region.

In this study, we uncovered a pathogenic role of meningeal immunity during neonatal meningitis, as well as its short- and long-term sequelae. We found that interleukin-17A (IL-17)-producing γδ T cells (γδ17 T cells) accumulate in the meninges of neonatal mice during group B streptococcal meningitis, and persist during adulthood. Importantly, our data show that meningeal γδ17 T cells contribute to blood-brain barrier (BBB) permeability during infection, facilitating bacterial invasion of the central nervous system (CNS) and microglia activation. This process is associated with attention deficit hyperactivity disorder (ADHD)-like behaviours in survivors. The uncovered mechanism underlying long-term behavioural sequelae of neonatal meningitis in mice opens the possibility of exploring new therapeutic approaches to alleviate brain damage in infants surviving group B streptococcal meningitis, increasing their quality of life.

## Results

### Clinical scoring does not predict neonatal death in the GBS meningitis model

We previously developed a mouse model of GBS neonatal meningitis in BALB/c mice^16^. Here, we adapted our model to the C57BL/6 strain by intravaginally inoculating pregnant females at gestational days (G) 16 and 17 (Figure 1a). We found that doses ranging from 3×10^4^ to 1×10^5^ colony-forming units (CFU) lead to neonatal disease, with dose-dependent mortality (Figure 1b). The best-fitting model, compared to our previous results obtained in the BALB/c model, was the colonisation with 3×10^4^ CFU, with half of the pups dying in the first five days of life (Figure 1b).

**Figure 1.**
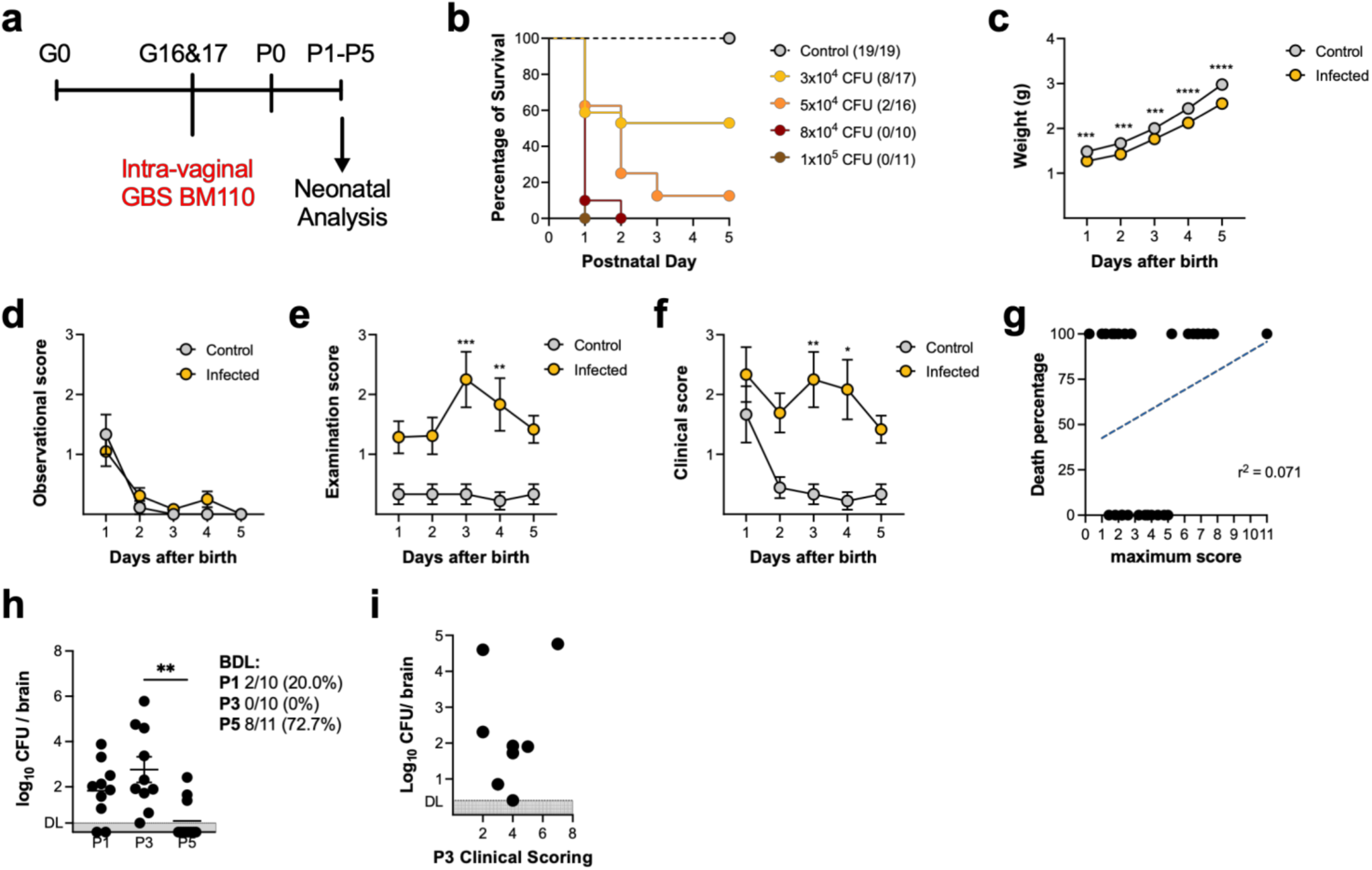
Clinical scoring does not predict neonatal death in experimental GBS meningitis. Pregnant C57BL/6 mice were intravaginally colonised with 10^4^ to 10^5^ CFU of GBS hypervirulent strain BM110, at gestational days 16 and 17. Control animals received PBS. **(a)** Schematic representation of the experimental protocol. **(b)** Kaplan-Meier survival curve of neonatal mice born from GBS-colonised dams, monitored for 5 days. The numbers in parentheses represent the number of animals that survived versus the total number of animals born. **(c)** Pups born from both groups were daily weighted until postnatal day (P) 5. Data is presented as mean values ± SEM (n = 9 Control, n = 21 Meningitis). Comparisons by Two-way ANOVA. **(d-f)** Observational **(d),** examination **(e)** and total **(f)** clinical scores of control and infected pups, during the first 5 days of life. Total clinical scoring was calculated by summing up the points from all the observational and examination categories. Data are presented as mean ± SEM (n = 9 Control, n = 21 Meningitis). Comparisons were performed by Two-way ANOVA. **(g)** Linear regression plot of the percentage of death vs the maximum clinical scores obtained from the infected pups. Each symbol indicates data from a single pup (n = 29). **(h)** Newborn mice were sacrificed at P1, 3, and 5 and the brain bacterial load was determined. Each symbol indicates data from a single pup (n = 10 P1, n = 10 P3, n = 11 P5). Data are presented as mean ± SEM. Comparisons by One-way ANOVA. **(i)** Correlation between the clinical scoring and the bacterial load in the brain of infected P3 pups. Each symbol indicates data from a single pup (n = 8). Statistical differences (P values) between groups are indicated. *P < 0.05, **P < 0.01, ***P < 0.001, ****P < 0.0001. DL, detection limit. BDL, below detection limit.

To assess disease progression and symptomatology appearance, as well as the predictability of death, in a non-invasive longitudinal and objectified sensitive manner, the offspring born from colonised and non-colonised dams were evaluated during their first five days of life for body weight and clinical scoring parameters (Figure 1c-g). As expected, infected animals presented significantly reduced body weight in all time points tested (Figure 1c). The observational score, composed of skin colour, presence of milk spot, spontaneous movement and head posture, showed no differences between groups (Figure 1d). In contrast, the examination score, comprising capillary refill time, dehydration, reaction to tactile stimuli, and abdominal palpation, was higher in the infected pups every assessed day, significantly worsening on postnatal days (P) 3 and 4 (Figure 1e). The total clinical scoring, combining observational and examination scores, was also significantly increased on these days (Figure 1f). Death predictability based on the clinical scoring was assessed by plotting the maximum obtained score *vs* the percentage of animals that died within the respective score (Figure 1g). The maximum score obtained varied between 1 and 11 (Figure 1g). Importantly, while a score equal to, or higher than, 6 predicted 100% mortality, pups with lower scores (score < 3) also exhibited high mortality (Figure 1g), limiting the scoring system’s predictive utility for death. Analysis of brain colonisation revealed that bacteria were already detected in 80% of the pups by P1, peaking at P3, and declining to nearly undetectable levels by P5 (Figure 1h). Noteworthy, brain bacterial burden did not correlate with clinical scores, as P3 animals exhibited variable brain colonisation at different clinical scores (Figure 1i), further emphasising the unpredictable nature of disease severity in this model. In conclusion, this model of neonatal meningitis does not rely on a direct correlation between bacterial levels in the brain, symptom appearance, and disease progression, unless a severe scenario of septicaemia is present.

### Neonatal GBS meningitis leads to an accumulation of meningeal **γδ** T cells

To define the meningeal immune landscape during neonatal meningitis, we dissected the dura mater and performed single-cell RNA sequencing (scRNA-seq) analysis on CD45^+^ immune cells sorted from pups with neonatal meningitis and uninfected controls at P3 (Figure 2a). The dura was chosen given its vast repertoire of immune cells^21^. This time point was selected as it corresponds to the peak of infection. To exclude possible confounders between cases of meningitis and septicaemia, pups with a clinical score higher than 6 were excluded. As a control for meningitis, brain colonisation was confirmed in all except one pup, as well as a significant decrease in brain weight compared to the control group (Figure 2b and c). After initial quality control, the generated dataset included 15766 cells from both control and infected mice. Cells were visualised using uniform manifold approximation and projection (UMAP), which revealed diverse immune subsets spanning across 21 distinct clusters (Figure 2d). The clusters were annotated based on curated published markers as well as data driven differentially gene expression analysis^32,33^ (Figure 2e-f). We identified multiple populations, including border-associated macrophages (BAM) (upregulating *Mrc1*, *Lyve1*, *Stab1* and *Dab2*), classical, intermediate and non classical monocytes (upregulating *Cd15*, *Ly6c2*, *Fcgr3*, *Eno3* and *Ace*), mast cells (upregulating *Tpsb2*, *Mcpt4* and *Mrgprb1*), basophils (upregulating *Cd200r3*, *Iffitm1*, *Ccl4*, *Ccl9* and *Aqp9*), neutrophils (upregulating *Ly6g*, *S100a9* and *Retnlg*), dendritic cells (DCs) (upregulating *H2-Ab1*, *Cd74* and *Flt3*), immature and mature B cells (upregulating *Cd19*, *Cd38* and *Pax5*, *Ms4a1*, *Cd79a*, *Ighm* respectively), apha beta and gamma delta T cells (upregulating *Trac*, *Trbc1* and *Trdc*, *Trgx1* respectively), a mixture of innate lymphoid cells (ILCs) and natural killer cells (NK) (upregulating *Zbtb16*, albeit at a lower level than γδ T cells, alongside the expression of *Klrb1c* and *Rorc* while lacking *Trac* and *Trdc* expression), contaminating osteoclast (upregulating *Acp5*, *Mmp9/14* and *S100a4*) and microglia (upregulating *Sall1*, *Sparc*, *Serpine2* and *P2ry12*), and a small cluster of fibroblast (upregulating *Col1a1* and *Col3a1*) (Figure 2e-f). BAMs appeared as the main cluster, followed by monocytes and mast cells, while lymphocyte clusters were much smaller (Figure 2g). The frequency of immune cell populations revealed differences between control and meningitis pups, including increased BAMs (Figure 2h). As expected, meningeal cells isolated from pups with meningitis revealed upregulation of genes related to the immune response to infection, such as *Isg15*, *Stat1*, *Fcgr4, C2* and *Cfh*, when compared to controls (Figure 2i). Based on these genes, the gene-set enrichment analysis (GSEA) of gene ontology (GO) terms showed that the total meningeal cells of meningitis pups are differentially activating pathways related to the immune activation, such as regulation of endocytosis, cell migration and motility, response to external stimuli, including bacteria and leucocyte and myeloid differentiation (Figure 2j). Comparing the ranking signature of the response to bacterium pathway within each cell type, we observed a marked increase in this pathway activity in the meningitis group across most meningeal immune populations, consistent with the infected state of these animals (Figure 2k). The response was most pronounced in neutrophils, monocytes, and BAMs, which showed the highest enrichment scores. Other immune subsets, including B cells, mast cells, γδ T cells and ILC/NK cells, also displayed elevated responses, albeit to a lesser extent (Figure 2k). Due to the low abundance neutrophils, classical mediators of the meningitis inflammatory response, and lymphoid cells, we further validated our findings by flow cytometry (Figure 3a and e, gating strategies). In agreement with their reduced body and brain weight (Figure 1c and 2c), meningitis pups displayed a significant decrease in the total number of meningeal immune cells at the peak of the infection (Figure 3b, kinetics shown in Figure S1a). No differences were observed either in CD11b^+^Ly6G^-^ macrophages/monocytes or in CD11b^+^Ly6G^+^ neutrophils in the dura during GBS meningitis (Figure 3 c and d, kinetics shown in Figure S1e and f). The frequency and number of B cells were significantly decreased in infected animals (Figure 3f, kinetics shown in Figure S1b). No differences were observed in αβ T cells (Figure 3g, kinetics shown in Figure S1c). Notably, the frequency of γδ T cells at P3 was significantly increased in the meningitis pups, almost doubling those from the control counterpart (Figure 3h, kinetics shown in Figure S1d). Systemically, increased proportions and numbers of γδ T cells were observed in the spleen of pups with meningitis at P1 and P3 (Figure S1j). At the peak of infection, the frequency of T cells was decreased in the spleen of infected animals, with the remaining populations not being affected (Figure S1g-l). Notably, no differences were observed in the main lymphoid and myeloid cells in the brain parenchyma (Figure S2), confirming our previous observation that GBS meningitis does not lead to leucocyte infiltration^16^.

**Figure 2.**
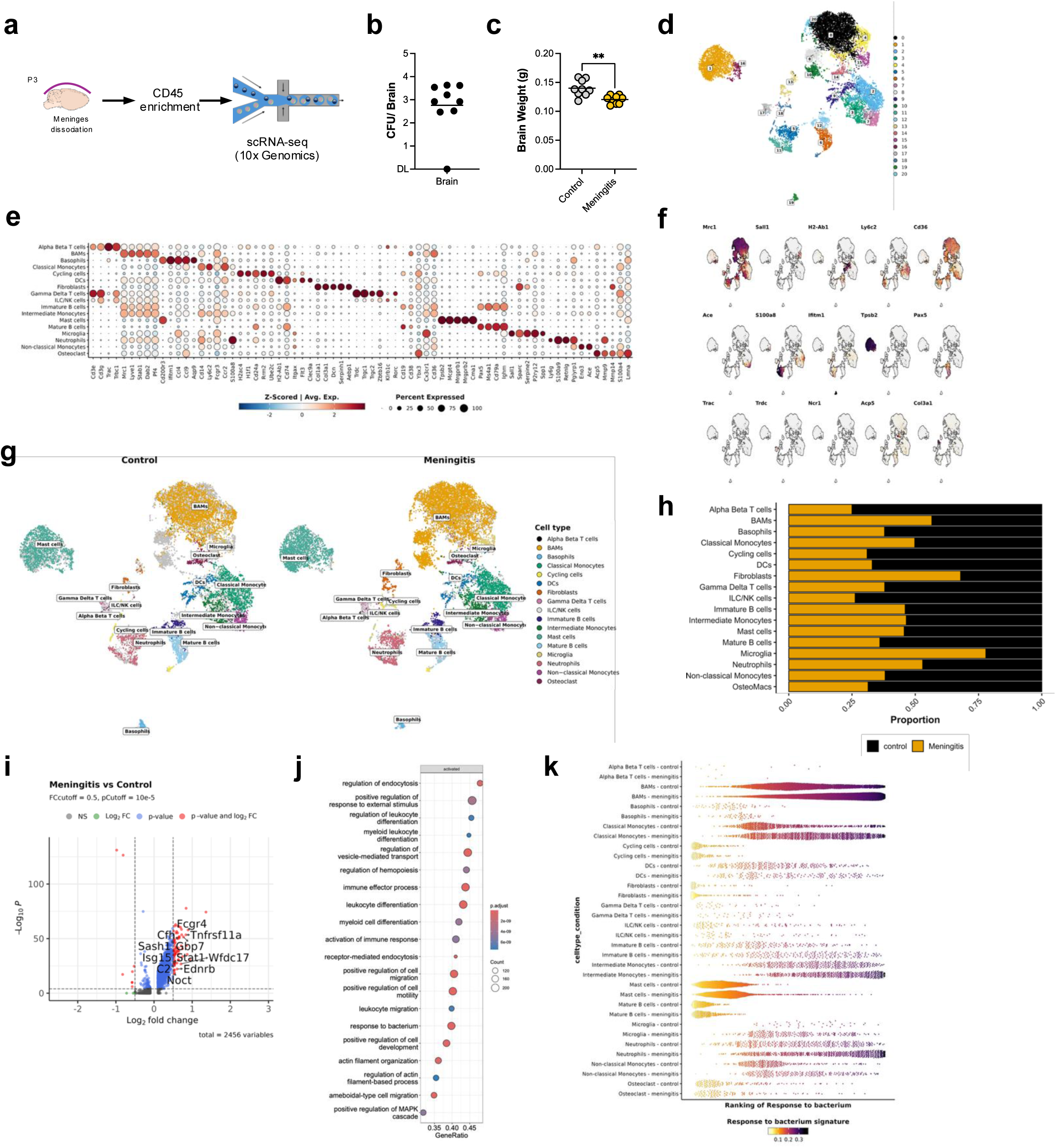
Composition of meningeal cell populations during neonatal meningitis by scRNA-seq. Pregnant C57BL/6 mice were intravaginally colonised with 3×10^4^ CFU of GBS hypervirulent strain BM110, at gestational days 16 and 17. Control animals received PBS. Newborn mice were sacrificed on postnatal day (P) 3. The meninges from 9 pups per group were harvested, pooled and processed for single-cell RNA sequencing (scRNA-seq). **(a)** Schematic illustration of P3 dural meninges isolation for scRNA-seq analysis. **(b-c)** The brain bacterial loads **(b)** and brain weight **(c)** were determined. Each symbol represents one pup (n = 9). Data are presented as mean. Comparison by Student’s t-test. **P < 0.01. DL, detection limit. **(d)** Uniform manifold approximation and projection (UMAP) of sorted dural meningeal cells cells (n = 15766 cells). **(e)** Dot plot showing Z-score for normalised average expression for selected publicly available curated markers for each indicated cell type. The colour and size of each dot represent the z-score and percentage of cells expressing each gene, respectively. **(f)** Feature plots of selected genes. The colour depicts the average gene expression. **(g)** UMAP of the distribution of CD45^+^ cell types in the dural meninges at steady state (Control) and during GBS meningitis. **(h)** Filled bar plot displaying the frequency of cell populations in Control vs Meningitis. **(i)** Volcano plot showing the differencially expressed genes in Meningitis vs Control. **(j)** Gene Set Enrichment Analysis (GSEA) of Gene Ontology (GO) terms showing the top 20 activated pathways by total cells in Meningitis vs Control. **(k)** Bee swarm plots for KNN smoothed UCell signature enrichment score of genes in GO response to bacterium pathway.

**Figure 3.**
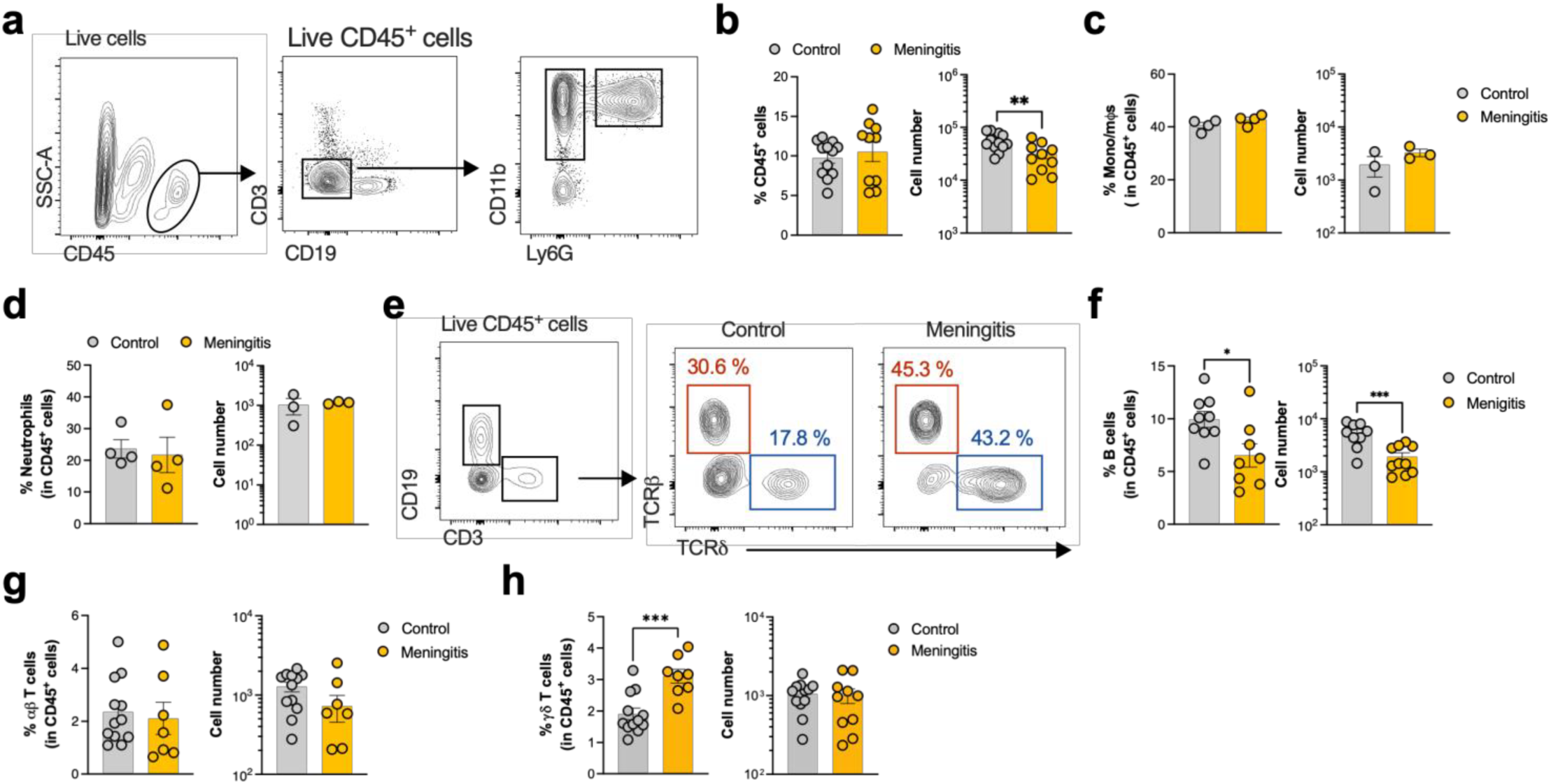
Increased frequency of γδ T cells in the meninges of GBS-infected newborn mice. Pregnant C57BL/6 mice were intravaginally colonised with 3×10^4^ CFU of GBS hypervirulent strain BM110, at gestational days 16 and 17. Control animals received PBS. Newborn mice were sacrificed at postnatal day (P) 3. The meninges were harvested and processed for flow cytometry analyses. **(a)** Representative flow cytometry scheme showing the myeloid gating strategy. **(b-d)** Frequency and number of indicated populations. Each symbol represents a single pup (n = 3 - 12). Error bars indicate the mean ± SEM. Comparisons by student’s t-test. **(e)** Representative flow cytometry scheme showing the lymphoid gating strategy. **(f-g)** Frequency and number of indicated populations. Each symbol represents a single pup (n = 12 Control, n = 10 Meningitis). Error bars indicate the mean ± SEM. Comparisons by student’s t-test. Statistical differences (P values) between groups are indicated; *P < 0.05, **P < 0.01, ***P < 0.001.

### **γδ**17 T cell accumulation upon neonatal meningitis persists in adulthood

It has been reported that male infants surviving invasive GBS disease are at higher risk of developing NDI^34^. Thus, we next evaluated whether sex could affect the frequency of meningeal γδ T cells at P3 (Figure 4a). These cells were found at higher proportions in the meningitis group, regardless of pup sex (Figure 4b). The same pattern was observed in the spleen, where infected animals exhibited an increased frequency of these cells regardless of sex (Figure S3a). Moreover, there were no sex-biased differences in bacterial levels in the brain and meninges (Figure 4c). Of note, the frequency of γδ T cells did not correlate with bacterial burden in the brain (Figure 4d). After confirming the increased frequency of γδ T cells in the meningitis group independent of sex, we sought to determine potential functional immune implications in this context. Under healthy conditions, meningeal γδ T cells are bona fide IL-17 producers (γδ17 T cells), with a T cell receptor (TCR) repertoire mostly restricted to the gamma chain variable region 6 (Vγ6) (assumed by default as double negative for Vγ1 and Vγ4)^26,29^. To assess whether neonatal meningitis leads to an altered phenotype of meningeal γδ T cells, we characterised this subset at P3. Flow cytometry analysis revealed that the TCR repertoire of γδ T cells during meningitis remained biased toward the Vγ6 subset (Figure 4a and e). Accordingly, intracellular staining showed that most meningeal γδ T cells in both groups remained IL-17 producers, with a negligible population of IFN-γ-producing γδ T cells (Figure 4a and f). Complementary scRNA-seq analyses of cytokine production, including IL-10, IL-6, IL-1β, TNF-α, and IL-17, revealed that γδ T cells are the major source of IL-17 in the meninges (Figure 4g), a result confirmed by flow cytometry (Figure 4h). Unexpectedly, we observed a significant decrease in the frequency of IL-17^+^ cells among γδ T cells in pups with bacterial meningitis (Figure 4f). Despite this, an overall increased IL-17 secretion by meningeal cells during meningitis was further validated by *ex vivo* stimulation with phorbol myristate acetate (PMA) and ionomycin (Figure 4i). IFN-γ levels were below the detection threshold, while measurable levels of IL-17 were only detected in the infected group, even without stimulation (Figure 4i). The GSEA analyses of the IL-17 signalling pathway showed that γδ T cells from meningitis pups also respond to IL-17 (Figure S3b). Notably, neutrophils, the first mediators of immune responses, also increase IL-17 signalling (Figure S3b). No differences were observed in the secretion of TNF-α, IL-6, or IL-10 (Figure S3c). Although no differences were found in the frequency of splenic IL-17-producing γδ T cells, accumulation of this cytokine was also found in culture supernatants from stimulated cells in the meningitis group (Figure S3d and e). γδ T cells progressively seed the meninges from early life and display a particularly activated tissue-resident phenotype^26,29,35^. GBS meningitis did not alter their phenotype, as Ki67 staining revealed that meningeal γδ T cells exhibited high proliferation at P3 (Figure S3f). In support of their activated resident phenotype, a fraction of meningeal γδ T cells expressed the tissue residency markers PD1 and CD69 in both control and meningitis groups (Figure S3g and h). Moreover, almost all γδ T cells displayed an activated CD44^+^CD62L^-/lo^ phenotype in both groups, showing no alterations in their tissue-homing and activation states (Figure S3i). CD44^+^CD62L^+^ cells, an effector memory phenotype, showed a tendency but not a significant increase in the meningitis group (Figure S3i).

**Figure 4.**
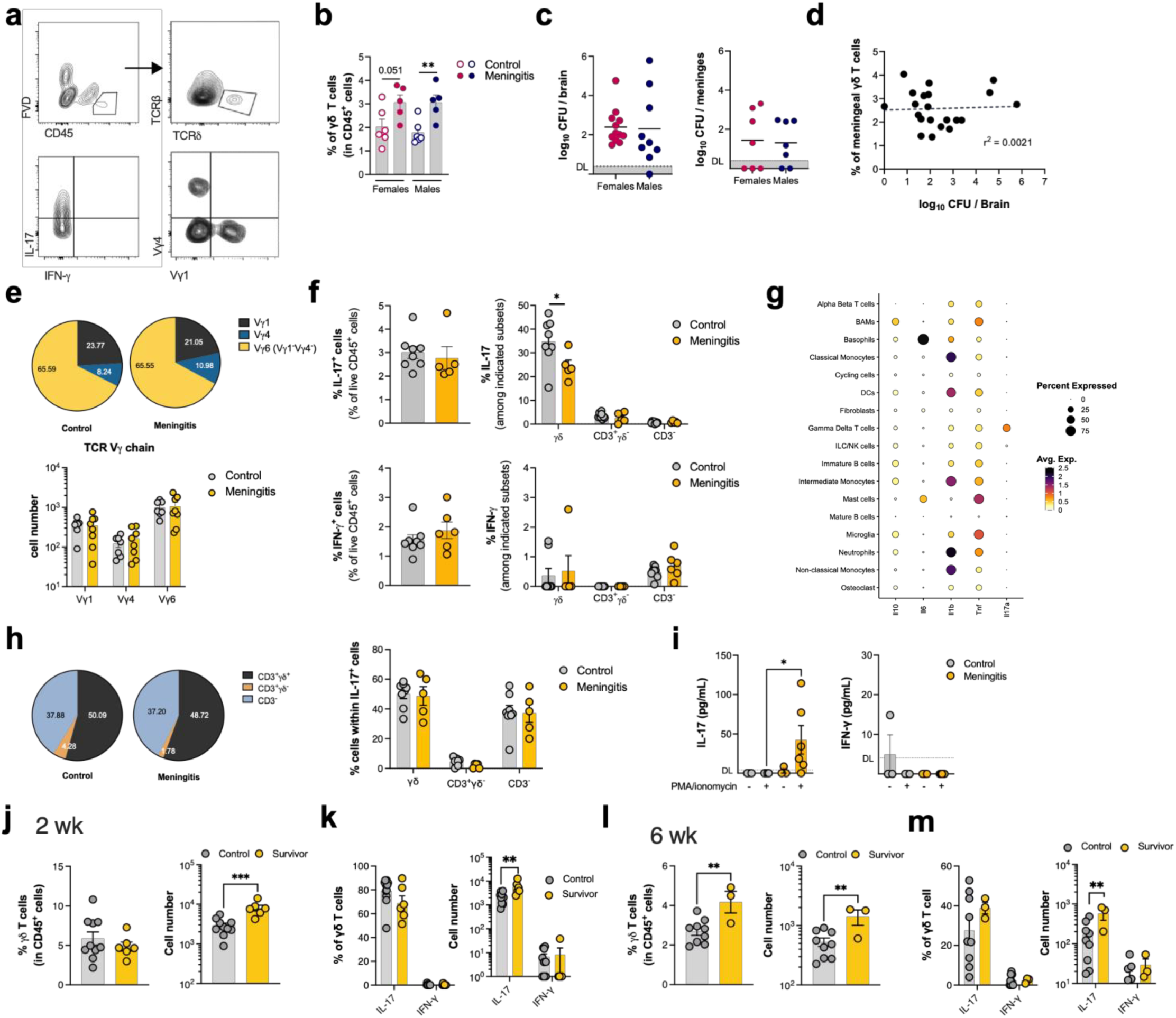
Neonatal meningitis alters γδ T cells IL-17 production in the meninges. Pregnant C57BL/6 mice were intravaginally colonised with 3×10^4^ CFU of GBS hypervirulent strain BM110, at gestational days 16 and 17. Control animals received PBS. **(a)** Representative gating strategy of the γδ T cells analysis. γδ T cells were defined as FVD^-^ CD45^+^ CD19^-^Tcrβ^-^TCRδ^+^. **(b)** Frequency of γδ T cells in the meninges of male and female pups at postnatal day (P) 3. Each symbol represents one pup (n = 6 Control, n = 5 Meningitis). Data are presented as mean ± SEM. Comparisons by Student’s t-test within sexes. **(c)** Bacterial load in the brain (left, n = 12 Females, n = 9 Males) and meninges (right, n = 7) in male and female pups, at P3. Each symbol represents one pup. Comparisons by Student’s t-test. **(d)** Linear regression plots of the frequency of γδ T cells and brain colonisation in P3 pups. Each symbol represents one newborn (n = 21). **(e)** Pie charts depicting the percentage of meningeal Vγ1^+^, Vγ4^+^ and Vγ6 (Vγ1^-^Vγ4^-^) γδ T cells, at P3 (top panel) and columns represent their frequency by animal (bottom panel). Each symbol represents one pup (n = 7 Control, n = 8 (Meningitis). Error bars indicate mean ± SEM. Comparisons by Student’s t-test within each clone. **(f)** Frequency of IL-17 (top panel) and IFN-γ (lower panel) produced by meningeal γδ T cells at P3. Each symbol represents one pup (n = 8 Control, n = 6 Meningitis). Error bars indicate mean ± SEM. Comparisons by Student’s t-test between groups. **(g)** Dot plot showing the scaled log-normalised average expression of selected cytokines. **(h)** Pie charts represent indicated cell populations within total IL17^+^CD45^+^ cells in the meninges (left panel), and column graphs represent the mean frequency of the indicated populations (right panel). Each symbol represents one pup (n = 8 Control, n = 6 Meningitis). Error bars indicate mean ± SEM. Comparisons by Student’s t-test within subsets. **(i)** Supernatant concentrations of indicated cytokines from *ex vivo* cultured meningeal cells, isolated from P3 pups, after 24 h of PMA and ionomycin stimulation. Each symbol represents a pool of 2 pups (n = 4-12). Data are presented as mean ± SEM. Comparisons by Student’s t-test within PMA absence or presence. **(j-m)** Frequency and number of γδ T cells and cytokines produced by them at 2 week-**(j-k)** and 6 week-**(l-m)** old control vs survivors. Each symbol represents one animal (n = 10 Control 2 wk, n = 9 Control 6 wk, n = 6 Meningitis 2 wk, n = 3 Meningitis 6 wk). Error bars indicate mean ± SEM. Comparisons by Student’s t-test. DL, detection limit.

Since γδ T cells have been shown to produce IL-17 in response to bacterial antigens and pro-inflammatory signals, such as IL-1β and IL-23^36^, we next tested whether GBS could directly modulate IL-17 production by these cells. To this end, γδ T cells were sorted from the neonatal lung, where they are highly enriched in the Vγ6^+^ subset (Figure S4), and stimulated with heat-killed GBS (hkGBS) in the presence or absence of splenocytes as a source of antigen-presenting cells (APCs) (Figure S3j). γδ T cells stimulated with hkGBS increased the production of IL-17, in the absence or presence of inflammatory stimuli (Figure S3j). Interestingly, APC presence potentiated cytokine production by γδ T cells (Figure S3j). To confirm that GBS induced the secretion of IL-1β and IL-23, splenic cells were stimulated with hkGBS alone. We found a significant increase in both cytokines, suggesting that *in vivo* GBS could further fuel γδ T cell activation (Figure S3k).

Finally, to evaluate whether γδ T cell homeostasis is restored following neonatal meningitis, flow cytometric analysis was performed on 2-week- and 6-week-old mice (Figure 4j-m). Surprisingly, the 2-week-old pups that have survived neonatal meningitis still presented bacterial colonisation in the brain, which was no longer detected at 6 weeks (Figure S5). Interestingly, meningeal γδ T cells remained significantly increased in meningitis survivors at both time points (Figure 4j and l). Intracellular cytokine staining confirmed the IL-17-production bias of survivors’ γδ T cells (Figure 4k and m). Altogether, these results show that GBS neonatal meningitis leads to γδ17 T cell accumulation in the meningeal microenvironment, persisting throughout life.

### **γδ** T cells drive GBS brain invasion in neonatal meningitis

To comprehend the contribution of γδ T cells to host protection against neonatal group b streptococcal meningitis, we next performed infection studies in wild-type (WT) and γδ T cell-deficient (*Tcrd^-/-^*) pups. Interestingly, *Tcrd^-/-^* infected pups were slightly more protected than WT pups (P = 0.091) (Figure 5a). Moreover, both groups showed no difference in bacterial loads in the lungs or meninges at P1 and P3 (Figure 5b and c). However, the *Tcrd^-/-^* group had lower bacterial burden in the brain at both time points, indicating that the absence of γδ T cells protects the brain from bacterial colonisation (Figure 5d). Analysis of immune cell populations in the meninges and spleen showed no differences in lymphoid (B cells and T cells) and myeloid (neutrophils, Ly6C^hi^ monocytes and macrophages) populations between infected and uninfected *Tcrd^-/-^* pups at P3 (Figure S6).

**Figure 5.**
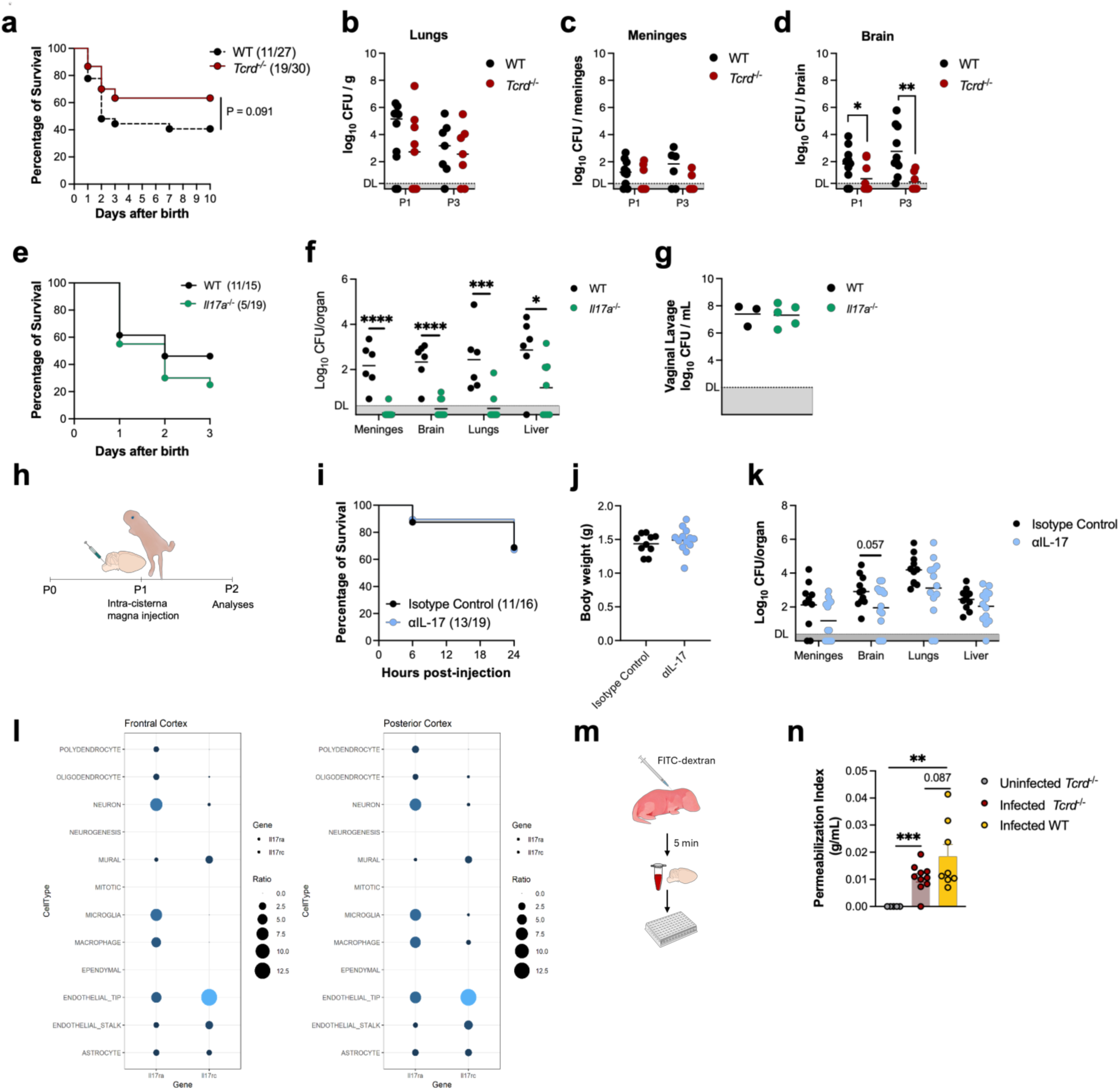
Meningeal γδ17 T cells increases susceptibility to GBS meningitis through BBB disruption. Pregnant C57BL/6 WT, *Il17a*^-/-^ or *Tcrd*^-/-^ mice were intravaginally colonised with 3×10^4^ CFU of GBS hypervirulent strain BM110, at gestational days 16 and 17. Control animals received PBS. **(a)** Kaplan-Meier survival curve of neonatal mice born from GBS-colonised WT and *Tcrd^-/-^* dams, monitored for 10 days. The numbers in parentheses represent the number of animals that survived versus the total number of animals born. **(b-d)** Bacterial load in the lungs **(b)**, meninges **(c)** and brain **(d)** at postnatal day (P) 1 and 3 in WT vs *Tcrd^-^*^/-^ pups. Each symbol represents one pup (n = 6-10 Control, n = 7-9 Meningitis). Error bars indicate mean ± SEM. Student’s t-test performed analysis. **(e)** Kaplan-Meier survival curve of neonatal mice born from GBS-colonised WT and I *Il17a*^-/-^ dams, monitored for 3 days. The numbers in parentheses represent the number of animals that survived versus the total number of animals born. **(f)** Bacterial load in meninges, brain, lungs and liver of postnatal (P) 3 WT and *Il17a*^-/-^ pups. Each symbol represents one pup (n = 6 WT, n = 9 *Il17a*^-/-^). Student’s t-test analysis. **(g)** GBS colonisation in vaginal lavage of vaginal colonised WT and *Il17a*^-/-^ dams one day after delivery. Each symbol represents one dam (n = 3 WT, n = 5 *Il17a*^-/-^). Comparison by Student’s t-test. **(h-k)** Anti-IL-17A or the Isotype IgG control was administered via intra-cisterna magna to WT P1-infected pups. Animals were analysed 24 hours postinjection (hpi). **(h)** Representative scheme of the αIL-17A administration protocol. **(i)** Kaplan-Meier survival curve of neonatal mice αIL-17A injected and controls, monitored for 24 h. The numbers in parentheses represent the number of animals that survived versus the total number of animals injected. **(j)** Body weight of αIL-17A and Isotype Control injected pups 24 hpi. Each symbol represents one pup (n = 10 Isotype Control, n = 13 αIL-17A). Student’s t-test analysis. **(k)** Bacterial load in meninges, brain, lungs and liver 24 hpi. Each symbol represents one pup (n = 11 Isotype Control, n = 13 αIL-17A). Student’s t-test performed analysis. **(l)** Predicted expression of the IL-17 receptor chains a (*Il17ra*) and c (*Il17rc*) in the different cell types, in the frontal (left plot) and posterior (right plot) cortex regions, using previously published single-cell RNA-seq data. The size of each dot represent the average expression of cells expressing each gene. **(m-n)** Postnatal day 2 pups born from were subcutaneously injected with FITC-dextran for BBB permeability evaluation. **(m)** Representative scheme of the blood-brain barrier (BBB) permeability assay. **(n)** Permeability index of the BBB of P2 uninfected *Tcrd^-^*^/-^ and infected WT and *Tcrd^-^*^/-^ pups. Each symbol represents one pup (n = 5 Uninfected, n = 10 Infected *Tcrd^-^*^/-^, n = 8 Infected WT). Student’s t-test analysis. Statistical differences (P values) between groups are indicated; *P < 0.05, **P < 0.01, ***P < 0.001, ****P < 0.0001. DL, detection limit.

As we found that meningeal γδ T cells are mostly IL-17 producers, we next sought to determine the role of this cytokine in the pathogenesis of GBS infection. No significant differences were observed in the survival of infected IL-17A-deficient (*Il17a*^-/-^) compared to WT pups (Figure 5e). However, *Il17a*^-/-^ pups exhibited a lower bacterial burden in the meninges, brain, lungs, and liver, with most having bacterial levels below the detection limit (Figure 5f). Although unlikely, *Il17a*^-/-^ progenitors could have increased control of GBS from their vaginal mucosa, leading to decreased vertical transmission. However, vaginal lavages performed one day after delivery confirmed comparable vaginal colonisation levels between *Il17a*^-/-^ and WT dams (Figure 5g). Collectively, these data suggest that IL-17 contributes to GBS pathogenesis rather than host protection in neonates.

We next asked whether locally targeting meningeal IL-17 could improve the outcomes of bacterial meningitis. Pups born from colonised progenitors were injected with IL-17-specific monoclonal antibody (αIL-17), or isotype control, directly into the cerebral spinal fluid (CSF, intra-cisterna magna, i.c.m.) at P1 and analysed 24 h later (Figure 5h). Local neutralisation of IL-17 did not improve neonatal survival (Figure 5i) or affect body weight (Figure 5j). However, while bacterial loads did not differ in the meninges or systemic organs (lungs and liver), αIL-17 injection directly into the pups’ CSF decreased bacterial load in the brain (P = 0.057) (Figure 5k), indicating local protection. Altogether, our data suggest that γδ T cells and IL-17 play a key role in facilitating bacterial colonisation of the brain during neonatal meningitis. To better understand the underlying mechanisms, we investigated the distribution of IL-17 receptor (IL-17R, both IL-17Ra and IL-17Rc), using available single-cell sequencing dataset^37^. IL-17R was expressed on several cell types, across different brain regions (Figure 5I and S7). Nevertheless, in regions proximal to the meninges, such as the cortex, endothelial cells were among the populations with a higher expression of IL-17R (Figure 5l). Based on these observations, we hypothesised that the blood-brain barrier (BBB) permeability might be differentially affected in infected *Tcrd*^-/-^ and WT pups. Thus, the integrity of the BBB was assessed *in vivo* at P3 using a FITC-dextran assay (Figure 5m). Uninfected *Tcrd*^-/-^ neonates were used as controls. Following systemic FITC-dextran injection, fluorescence levels in the blood and the brain were quantified. Both infected WT and *Tcrd*^-/-^ pups showed significant BBB permeability compared to uninfected *Tcrd*^-/-^ pups (Figure 5n). Importantly, higher permeability was detected in the infected WT than in *Tcrd*^-/-^ neonates (P = 0.087), indicating that the absence of γδ T cells helped maintain BBB integrity. In summary, these findings show that meningeal γδ T cells are important mediators of neonatal bacterial meningitis, facilitating CNS infection through IL-17 secretion and BBB disruption.

### Meningeal **γδ**17 T cells in neonatal meningitis leads to reminiscent of ADHD-like behaviour

Given that infants surviving neonatal meningitis experience permanent NDI and that meningeal immunity influences CNS function^16,38,39^, we investigated whether γδ T cells contribute to altered behaviour phenotypes in this model. Behavioural assessments were performed throughout the lactation period by evaluating the acquisition of developmental milestones, and later during middle adolescence (P42), using a battery of classical behaviour tests. Both male and female animals were included during the developmental phase to provide a comprehensive understanding of early-life effects across sexes and mimicking natural litter compositions. The developing reflex of audition revealed that infected pups, regardless of the presence of γδ T cells, anticipate this sense by approximately 2 days compared to their respective controls (Figure 6a). The ear twitch, a reflex involving ear movement in response to touch, showed differences between genotypes (Figure 6a). While infected WT pups presented earlier reflex compared to uninfected control ones, there was no difference between *Tcrd*^-/-^ groups (Figure 6a). No differences were observed in surface righting among groups (Figure 6b). Eye opening occurred significantly earlier in the infected WT neonates, whereas no differences were found between the *Tcrd*^-/-^ groups, similarly to the ear twitch reflex results (Figure 6c). Regarding strength and coordination, both infected WT and *Tcrd*^-/-^ neonates presented delayed cliff aversion reflex (Figure 6d). Anterior member strength was assessed through the forelimb grasp test, revealing that infected WT animals completed the test 1.5 days earlier than the control group, similarly to what was observed in both the control and infected *Tcrd*^-/-^ pups (Figure 6d). No differences were found in air righting among groups (Figure 6d). Interestingly, locomotion assessment revealed that infected WT pups completed the open field traversal test in 5 seconds, half of the time needed by control pups (Figure 6e), suggesting hyperactivity in the meningitis group. No differences were found between the *Tcrd*^-/-^ groups (Figure 6e).

**Figure 6.**
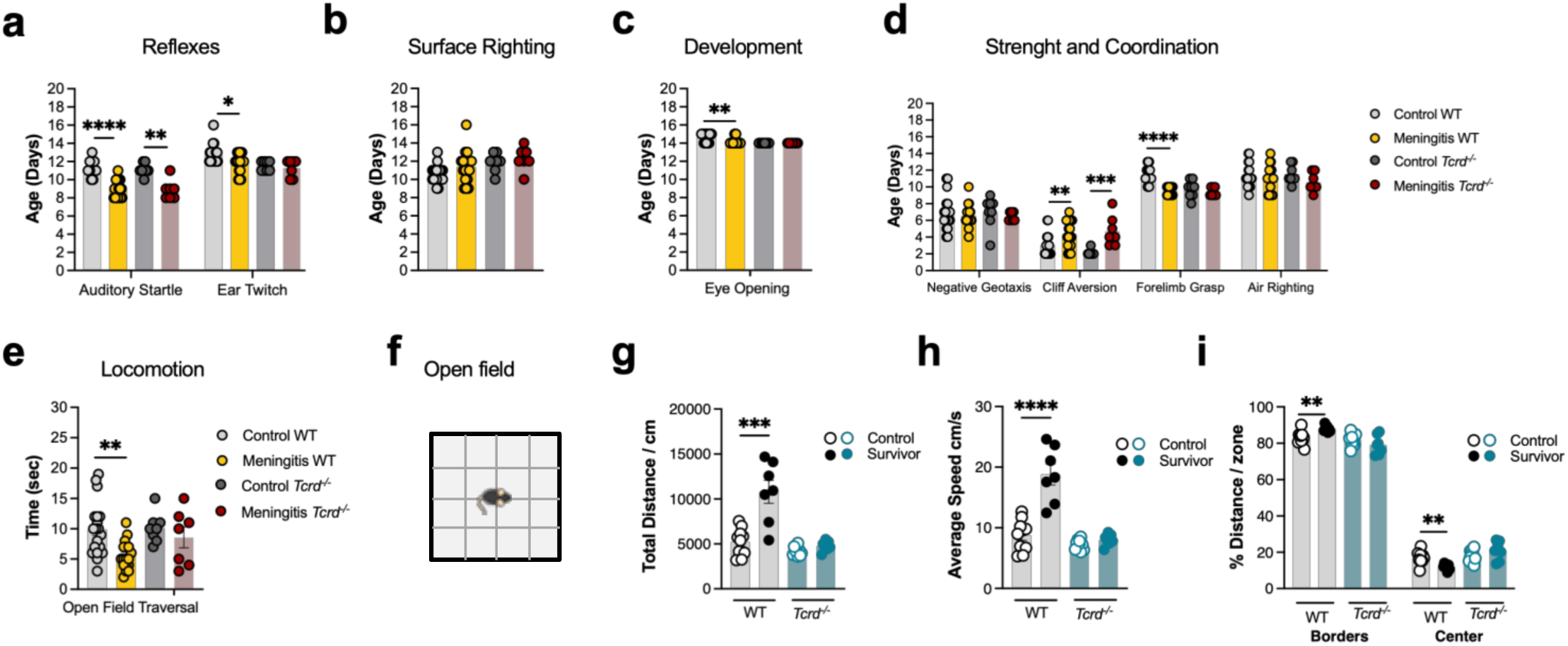
The absence of γδ T cells restores developmental milestones and behaviour in meningitis survivors. Pregnant C57BL/6 WT or Tcrd^-/-^ mice were intravaginally colonised with 3×10^4^ CFU of GBS hypervirulent strain BM110, at gestational days 16 and 17. Control animals received PBS. **(a-e)** Pups born from the indicated groups were tested daily for the first 21 days of life for the indicated reflexes and developmental signs, with the tasks determined at specific time points according to milestones. Each symbol indicates data from a single pup (n = 20 Control WT, n = 18 Meningitis WT, n = 8 Control *Tcrd^-/-^*, n = 7 Meningitis *Tcrd^-/-^*). Comparisons within genotypes, through Student’s t-test. **(f-i)** Male animals born from the indicated groups were allowed to reach adulthood and submitted to behavioural tests. **(f)** Representative scheme of the Open Field test. The total distance covered **(g)**, the average speed **(h)** and the percentage of distance covered by zone **(i)** were used as a measure of hyperactivity and anxiety-like behaviour. Each symbol represents a single animal (n = 10 Control WT, n = 7 Survivor WT, n = 9 Control *Tcrd^-/-^,* n = 7 Survivor *Tcrd^-^/^-^*). Error bars indicate mean ± SEM. Comparisons by Student’s t-test within genotypes and type of object. Statistical differences (P values) between groups are indicated; *P < 0.05, **P < 0.01, ***P < 0.001, ****P < 0.0001.

Given the known sex differences in adult behaviour and physiology^40^, males and females were assessed separately during adulthood (Figure 6, S8 and S9). Surviving WT female mice showed no altered behavioural phenotype, including locomotion, anxious behaviour, cognition and social preferences (Figure S8). When assessing males, no differences were found between groups and genotypes in the elevated plus maze test and sociability assessment (Figure S9a-g). Regarding cognition, through the novel object recognition test, survivors from both genotypes failed to recognise the novel object as being new, unlike their respective controls, showing impaired recognition memory regardless of γδ T cells’ presence (Figure S9h-k). However, in the open field (OF) test, surviving WT male mice exhibited significantly increased total distance travelled and higher average speed, indicating hyperactivity, a typical ADHD-like behaviour (Figure 6f-i). In addition, these animals also showed signs of open space anxiety, as evidenced by significantly greater distance at the borders of the OF arena with concomitant lower distance at the centre, despite no differences in the time spent/per zone (Figure 6i and Figure S9e). Notably, no differences were found in the absence of γδ T cells, with surviving *Tcrd*^-/-^ pups behaving as controls (Figure 6f-I and S9e). Together, these data indicate a preferential impact of γδ T cells on ADHD-like behaviours, particularly hyperactivity and anxiety, in the context of neonatal meningitis.

### Neonatal GBS meningitis leads to microglia activation

Having shown that meningeal γδ T cells facilitate bacterial entry into the CNS during GBS meningitis, contributing to ADHD-like behaviour, we further explored the mechanism underlying long-term sequelae of neonatal meningitis through bulk RNA-sequencing (RNA-seq) of the meninges from infected pups and healthy controls (Figure S10a). Principal component analysis (PCA) of all expressed genes showed a clear separation between uninfected pups and those with neonatal meningitis (Figure S10b). Differential gene expression analysis identified 569 genes that were upregulated in neonatal meningitis compared to healthy pups (Figure S10c). To identify gene sets with similar biological activity with significant changes in transcriptomic levels, we employed a gene set enrichment analysis (GSEA) using mouse hallmark gene sets (Figure S10d and e). Interestingly, all biological processes related to the immune response were found to be significantly downregulated during neonatal meningitis (Figure S10d). In contrast, cellular processes associated with development, differentiation and commitment revealed a significant increase in pathways associated with glia and neurons (Figure S10e). Genes having the highest variation within the top 7 up-regulated pathways are highlighted in Figure S10f.

Given the observed altered neuron and glia signature, we quantified the number of neurons and microglia, and assessed their reactivity in several relevant brain regions, namely the prefrontal cortex, somatosensory cortex and hippocampus, in male offspring at P3. Interestingly, in meningitis pups, decreased mean intensity of neuronal marker NeuN was observed in the hippocampus, indicating decreased numbers of neurons or a delay in neuron maturation (Figure 7a and b), while the microglia compartment, characterised by Iba-1, appeared unchanged in all tested brain regions (Figure 7c). We subsequently analysed the morphology of these cells, a commonly used parameter to assess the inflammatory phenotype^41^. Imaris semi-automatic 3D reconstructions of microglia revealed no differences in microglia surface area and volume between groups (Figure 7a, d and e). However, phagocytic activity, as determined by CD68 immunostaining, was significantly increased in the hippocampal microglia of meningitis pups, with a similar trend observed in the somatosensory cortex (P= 0.052) (Figure 7f and g). For deeper characterisation of the microglia status, we next performed flow cytometry analyses of the whole brain at P3 (Figure 7h and i). While the frequency of microglia was significantly increased in pups with meningitis, their number was statistically decreased, consistent with the decreased brain weight observed in this group at this time point (Figure 7h and 2c). Imaging data were corroborated by a significant increase in SSC-A mean fluorescence intensity (MFI), indicative of higher intracellular granularity and activity (Figure 7i). Moreover, all the other activation markers studied, including CD45, CD11b, F4/80 and CX3CR1, were significantly increased in the microglia of pups with meningitis, confirming their activation status as observed through CD68 quantification (Figure 7g and i). These results align with the active infection status in the parenchyma of these pups (Figure 1h). Of note, confocal microscopy analysis of adult brains from meningitis survivor and sham-infected mice showed no differences in microglia numbers and morphology, as well as no differences in neuronal quantification (Figure S11). Altogether, our data show that neonatal microglia are overall decreased during GBS meningitis, but are highly activated, consistent with the infection status of the animals.

**Figure 7.**
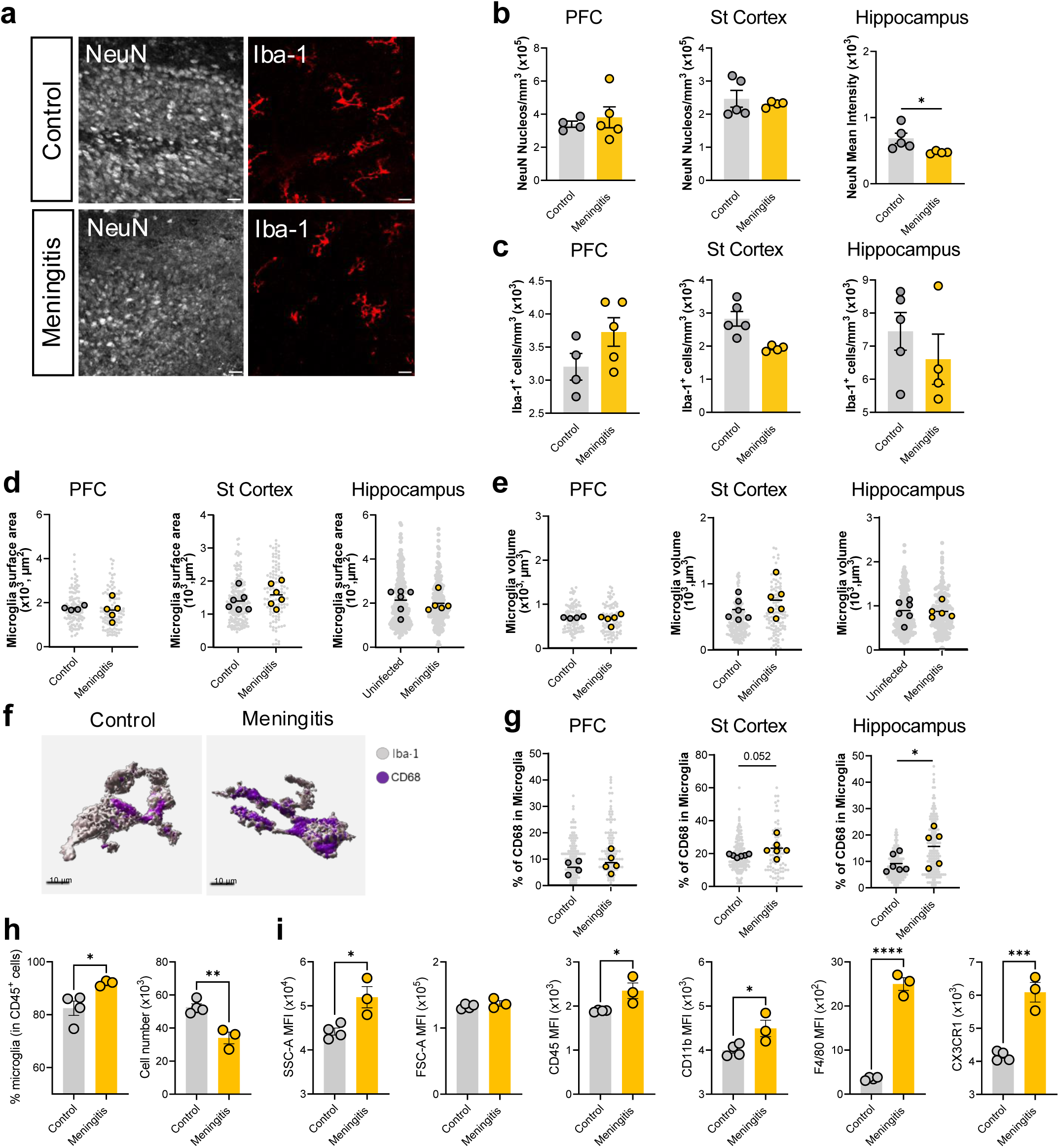
Neonatal GBS infection leads to increased microglial activation. Pregnant C57BL/6 mice were intravaginally inoculated with 3×10^4^ CFU of GBS hypervirulent strain BM110 at gestational days 16 and 17. Control animals received PBS. The male progeny was sacrificed at postnatal day (P) 3. **(a)** Representative images of NeuN and Iba1 in the cortex of P3 pups from both groups. Scale bar 20 μm. **(b)** Quantification of NeuN cells in the brain’s prefrontal cortex (PFC), somatosensory cortex (St cortex) and hippocampus. Each symbol represents one animal (n = 4-5 per group). Error bars indicate mean ± SEM. Comparisons by Student’s t-test. **(c)** Quantification of Iba-1 cells in the brain’s prefrontal cortex (PFC), somatosensory cortex (St cortex) and hippocampus. Each symbol represents one animal (n = 4-6 per group). Error bars indicate mean ± SEM. Comparisons by Student’s t-test. Microglia surface area **(d)** and volume **(e)** at the indicated brain regions. **(f-g)** Quantitative analysis of immunostaining for CD68 inside microglia in the indicated brain regions. Comparisons by Student’s t-test. **(f)** Representative Imaris 3D reconstruction of microglia labelled with Iba-1, from the cortex of the indicated groups. Lysosomes (CD68, purple) can be visualised inside microglia and quantified. **(g)** Quantification of CD68 in the indicated brain regions. Each coloured symbol represents one animal (n = 4-6) and each background grey symbol represents one cell (n = 115-257 Control, n = 106-175 Meningitis). Comparison by Nested t-test. **(h)** Frequency and number of microglia by flow cytometry. **(i)** Quantification of SSC-A, FSC-A, CD45, CD11b, F4/80 and CX3CR1 on microglia, presented as median fluorescence intensity (MFI). Each symbol represents one animal (n = 4 Control, n = 3 Meningitis). Error bars indicate mean ± SEM. Comparisons by Student’s t-test. Statistical differences (P values) between groups are indicated. *P < 0.05,**P < 0.01, ***P < 0.001, ****P < 0.0001.

### Meningeal **γδ**17 T cells associates with GBS virulence

Given that the absence of γδ T cells results in decreased bacterial load in the brain and diminished long-term sequelae, we next performed a detailed characterisation of microglial morphology and functional status in *Tcrd*^-/-^ pups. While Iba-1⁺ microglial cell numbers remained unchanged (Figure 8a), we observed a significant reduction in microglial surface area and volume in the somatosensory cortex, and no morphological alterations in the hippocampus (Figure 8b and c). Despite this, CD68 expression was significantly increased in microglia from the same regions (Figure 8d), indicating phagocytosis. The combination of reduced morphological complexity and increased lysosomal content may reflect a transition to a compact amoeboid phenotype, consistent with localised activation. Indeed, flow cytometric profiling of microglia from whole brains of *Tcrd*^⁻/⁻^ neonates revealed no significant changes in granularity (SSC-A), cellular complexity (FSC-A), or the expression of canonical activation markers, including CD45, CD11b, F4/80, and CX3CR1 (Figure 8e), suggesting that microglia in *Tcrd*⁻^/^⁻ pups mount a focally restricted and potentially more controlled response.

**Figure 8.**
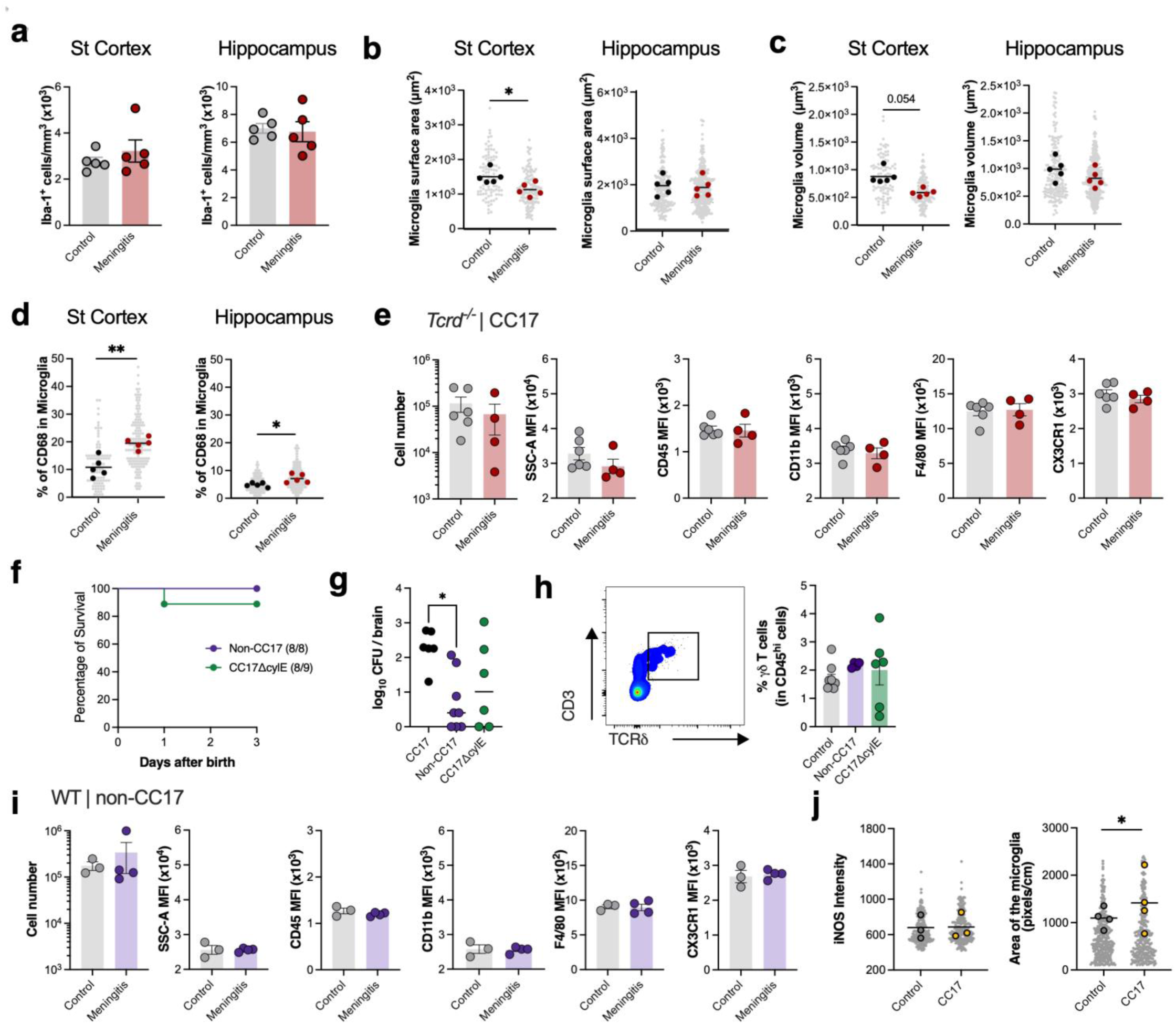
Microglia activation is associated with brain parenchyma invasion. **(a-e)** Pregnant *Tcrd^-/-^* mice were intravaginally inoculated with 3×10^4^ CFU of GBS hypervirulent strain BM110 at gestational days 16 and 17. Control animals received PBS. The male progeny was sacrificed at postnatal day (P) 3. **(a)** Quantification of Iba-1 cells in the brain’s somatosensory cortex (St cortex) and hippocampus. Each symbol represents one animal (n = 5). Error bars indicate mean ± SEM. Comparisons by Students’ t-test. Microglia surface area **(b)** and volume **(c)** at the indicated brain regions. Each symbol represents one animal (n = 5). Error bars indicate mean ± SEM. Comparisons by Students’ t-test. **(d)** Quantification of CD68 in the indicated brain regions. Each coloured symbol represents one animal (n = 5) and each background grey symbol represents one cell (n = 10-48 Control, n = 23-75 Meningitis). Comparison by Nested t-test. **(e)** Number of microglia by flow cytometry and quantification of SSC-A, CD45, CD11b, F4/80 and CX3CR1 on microglia, presented as median fluorescence intensity (MFI). Each symbol represents one animal (n = 6 Control, n = 4 Meningitis). Error bars indicate mean ± SEM. Comparisons by Students’ t-test. **(f-i)** Pregnant C57BL/6 WT mice were intravaginally inoculated with 3×10^4^ CFU of the indicated GBS strain at gestational days 16 and 17. Control animals received PBS. The male progeny was sacrificed at P3. **(f)** Kaplan-Meier survival curve of neonatal mice born from Non-CC17 and CC17ΔcylE GBS-colonised WT progenitors, monitored for 3 days. The numbers in parentheses represent the number of animals that survived versus the total number of animals born. **(g)** Bacterial load in the brain of infected pups at P3. Each symbol represents one pup (n = 6-8). Comparisons by One-way ANOVA. **(h)** Representative flow cytometry scheme showing the gate on γδ T cells (left) and the frequency of this population in Control, Non-CC17 infected and CC17ΔcylE infected at P3. Each symbol represents one pup (n = 9 Control, n = 4 Non-CC17, n = 6 CC17ΔcylE). Error bars indicate mean ± SEM. Comparisons by One-way ANOVA. **(i)** Number of microglia by flow cytometry and quantification of SSC-A, CD45, CD11b, F4/80 and CX3CR1 on microglia, presented as MFI. Each symbol represents one animal (n = 3 Control, n = 4 Meningitis). Comparisons by Student’s t-test. **(j)** Primary microglial cultures were stimulated with media (Control) or hkGBS (CC17) at a multiplicity of infection of 10 for 24 h. iNOS (left) by immunohistochemistry and microglia area (right). Data presented as mean. Each coloured symbol represents one culture (n = 3) and each background grey symbol represents one cell (n = 216-330 Control, n = 183-280 CC17). Comparisons by Nested t-test. Statistical differences (P values) between groups are indicated. *P < 0.05,**P < 0.01.

Taken together, these data suggest that γδ T cell activation and subsequent GBS invasion into the CNS are associated with enhanced microglial activation and neuronal injury. To test this hypothesis, we next performed infection studies in WT pups using bacterial strains less associated with meningitis, namely the serotype III NEM316 (non-CC17), which does not belong to hypervirulent clonal complex 17, and the attenuated CC17ΔcylE strain lacking haemolytic/cytolytic activities. Maternal colonisation with either strain resulted in high neonatal survival rates (Figure 8f). In accordance, both attenuated strains exhibited reduced bacterial burden in the brain compared to pups infected with the hypervirulent CC17 strain, reaching significance in the non-CC17 group (Figure 8g). We next assessed meningeal γδ T cells to determine whether their accumulation was affected by infection with less virulent strains. At P3, no differences were observed between the experimental groups (Figure 8h). We then examined whether pups infected with a less virulent strain exhibited reduced microglial reactivity. Flow cytometric analyses showed no alterations in microglia numbers or the expression of activation markers in pups infected with the non-CC17 strain (Figure 8i). Finally, to further explore whether the presence of bacteria is sufficient to activate microglia, primary microglia cultures were stimulated with hkGBS CC17 for 24 h. The results show that while hkGBS stimulation did not lead to increased expression of the classical microglial activation marker iNOS, it significantly increased microglial cell area, compared to control, indicative of activation (Figure 8j). Altogether, these results suggest that γδ T cells contribute to CNS invasion and GBS virulence by modulating microglial reactivity, highlighting their potential role in disease pathophysiology.

## Discussion

Neonatal meningitis remains a leading infectious cause of mortality and long-term disability globally^42^. However, the mechanisms leading to neurological sequelae are poorly understood. Here, we uncover a novel mechanism for NDI in GBS meningitis by demonstrating that bacterial stimuli drive lifelong accumulation of γδ17 T cells in the meninges. This process disrupts BBB integrity and activates microglia, leading to phenotypes resembling ADHD in mice.

Body weight loss during the first postnatal days of GBS infected pups confirmed systemic impact of infection, yet clinical scoring failed to predict mortality, emphasising the subtle and unpredictable nature of GBS meningitis. This mirrors the human disease, in which symptoms are often absent or nonspecific, complicating early diagnosis^43,44^. Notably, bacterial colonisation of the brain occurred rapidly after birth, but disease severity was not proportional to bacterial load, suggesting that host factors also contribute to the outcome.

The meninges contain a dense network of immune cells that intimately interact with the CNS, playing crucial roles in (patho)physiology^39,45^. Yet, their contribution to early-life immunity remains largely unexplored. Single-cell RNA-seq uncover a complex meningeal immune landscape already established at P3, closely resembling that of adults. Flow cytometry confirmed a significant increase of γδ T cells in infected pups, independent of bacterial burden in the brain, indicating that mere pathogen presence is sufficient to activate meningeal immunity. These γδ T cells retained their canonical phenotype^26,29^, including TCR Vγ6, tissue-residency markers, and a bias toward IL-17 production. Interestingly, although IL-17 levels were elevated in the meningeal microenvironment, transcriptional regulation of IL-17 in γδ T cells was dampened during meningitis. This aligns with previous findings that commensal-derived cues modulate meningeal γδ17 T cells^29^. Given that GBS colonises the neonatal gut^16,46^, it is plausible that the gut-meningeal axis can be modulating meningeal γδ17 T cells during infection.

Our data may have translational value for human neonates, as γδ17 T cells are among the earliest T cells generated in the foetal human thymus^47^ and have been shown to produce IL-17 in children with bacterial meningitis^48^. Importantly, the number of γδT17 cells in these children decreases after treatment, suggesting a pathogen-specific response^48^, as we also found here with γδ T cells directly responding to GBS. Studies with cord blood cells also showed that human neonatal γδ T cells can produce IL-17 upon polyclonal activation^49^. Additionally, γδ T cells have already been detected in the blood of extremely premature babies and the brain lesions of premature infants with white matter injury^50,51^. To assess whether the alterations in γδ T cells persisted beyond the neonatal period, we examined older survivors. IL-17-producing γδ T cells remained significantly elevated in these mice until adulthood, suggesting either increased proliferation or recruitment to the meninges. In steady-state, γδ T cells populate peripheral tissues during the perinatal period, where they proliferate and self-renew throughout life^52^. Although we did not find differences in proliferation, the intrinsically high turnover of γδ T cells under homeostatic conditions in the postnatal period may mask subtle changes. Alternatively, recruitment from peripheral pools cannot be excluded, as splenic γδ T cells were also increased at P3, and in the lungs of pups vertically infected with GBS^53^.

Although γδ T cells are traditionally associated with pathogen clearance and tissue repair^54^, their absence did not significantly improve survival in GBS-infected pups. Nevertheless, despite comparable bacterial colonisation in peripheral organs and meninges, *Tcrd*^-/-^ pups exhibited a marked reduction in bacterial invasion of brain parenchyma, suggesting a pathogenic role of meningeal γδ T cells to CNS entry. This phenotype was recapitulated in IL-17-deficient animals, which displayed reduced bacterial load across all tested tissues, indicating that IL-17, typically protective in other infectious settings, disrupts host-pathogen balance in neonatal GBS meningitis. While IL-17 is classically protective against extracellular bacteria through the recruitment of neutrophils^55^, in neonatal GBS meningitis it disturbs the tight balance between infection and inflammation. Rather than enhancing resistance, excessive IL-17 signalling amplifies inflammatory injury, potentially compromising BBB integrity and worsening pathology without improving bacterial clearance. In line with this, local IL-17 neutralisation in the cisterna magna reduced bacterial colonisation in the brain, further implicating IL-17 in BBB modulation. Consistently, *Tcrd*^-/-^ pups exhibited reduced BBB permeability, suggesting that meningeal γδ17 T cells may influence barrier integrity through local cytokine production. However, direct IL-17-mediated BBB disruption remains to be formally demonstrated, and other IL-17-producing populations, such as type 3 innate lymphoid cells (ILC3s), may contribute to this effect. Moreover, GBS has virulence factors capable of promoting CNS invasion independently of BBB breakdown^9,56–58^, which may explain residual bacterial colonisation observed in *Tcrd*^-/-^ pups. γδ17 T cells have already been implicated in other neurological diseases, including experimental autoimmune encephalomyelitis (EAE) and stroke^59^, where IL-17–driven neutrophil recruitment disrupts the BBB^60–65^. Our findings suggest a distinct mechanism in neonatal meningitis, in which meningeal γδ17 T cells modulate BBB integrity through local IL-17 release, emphasising the importance of inter-tissue communication in early-life CNS immunity. Together, these data highlight a fragile balance between infection control and inflammation, positioning IL-17 as a critical regulator of disease tolerance. In this context, IL-17 acts as a detrimental amplifier of inflammation, and its inhibition may restore homeostatic balance by limiting infection-induced injury in the developing brain. Finally, it should be acknowledged that no current model allows the selective depletion of IL-17 within γδ T cells, representing a limitation in dissecting their specific contribution to BBB disruption. Nevertheless, the downstream consequences of this meningeal inflammatory axis are evident within the brain parenchyma, where altered microglial activation becomes a key effector of neuronal injury.

Microglia from infected WT neonates exhibited a transcriptional and morphological profile consistent with reactive activation, whereas those from *Tcrd^-/-^* pups displayed a focally restricted and potentially more controlled response. This suggests that γδ17 T cells and IL-17 signalling amplify microglial reactivity, shifting it from a homeostatic to a disease promoter mode. Consistent with this, microglia activation correlated with the bacterial strain’s ability to invade the CNS, linking meningeal inflammation to parenchymal immune activation. Collectively, these data support a mechanism in which IL-17-mediated modulation of the BBB promotes bacterial entry, thereby enhancing microglial activation and neuronal injury.

Beyond acute pathology, the neuroimmune perturbations established in early life translated into persistent behavioural alterations. Accelerated sensory development and hyperactivity in WT survivors indicate aberrant neurodevelopmental trajectories reminiscent of attention-deficit and hypersensitivity phenotypes^66^. A rat model of maternal separation showed an acceleration of eye opening^67^, demonstrating that, similarly to our findings, early-life stress can precipitate premature sensory maturation. Likewise, neonatal infection has been shown to exacerbate the acoustic startle response in adulthood, but only when animals were first exposed to stress^68^, consistent with our observation of anticipatory auditory reflexes in infected pups. When reaching early adulthood, both WT and *Tcrd*^-/-^ male survivors presented deficits in recognition memory. This impairment, therefore, appears unrelated to γδ T cells or bacterial burden, suggesting additional mechanisms underlying learning and memory dysfunction in GBS meningitis. Future studies are needed to address this. In the open field arena, WT survivors exhibited hyperactive behaviour, as evidenced by higher total distance travelled and increased speed, particularly along the borders of the arena, compared to the control group. Their preference for the periphery and reduced travel distance in the center suggests that anxiety-related behaviour may coexist with increased activity levels. Notably, these behavioural changes were abrogated in the absence of γδ T cells, implicating a potential role for these immune cells in modulating long-term neurobehavioral outcomes. This is consistent with a previous study showing that γδ17 T cells seed the meningeal spaces and control anxiety-like behaviour in mice^29^.

Although anxiety-like behaviour was not confirmed in the EPM, the open field paradigm is particularly sensitive to hyperactivity and exploratory drive, suggesting that γδ17 T cells may preferentially modulate locomotor and attention-related circuits. These findings align with prior evidence implicating meningeal γδ17 T cells in regulating neuronal signalling, behaviour, and cognition in both homeostatic and pathological states^26,29,64,69,70^. Hyperactivity and anxiety frequently coexist, including in paediatric ADHD, which affects 5% of children worldwide, with 30-60% maintaining symptoms in adulthood ^71–74^. Future studies employing complementary paradigms, such as the light– dark box, novelty-suppressed feeding test, or long-term open-field assays, could help disentangle the components of anxiety versus hyperactivity. Comparable outcomes have been reported in models of maternal immune activation, where early immune challenges impair brain development and induce ADHD-like behaviours^75–77^. Moreover, human studies show that perinatal inflammation increases the risk of ADHD in the offspring^78–80^, though mechanistic links between early-life infection and disease onset remain scarce.

By combining scRNA-seq, flow cytometry, imaging, and behavioural assays with a clinically relevant model of neonatal meningitis, we demonstrate a causal link between meningeal γδ17 T cells dysregulation, bacterial invasion into the CNS, and subsequent permanent neurodevelopmental deficits, including ADHD-like behaviour. This study uncovers a previously unrecognised neuroimmune pathway in GBS meningitis and suggests IL-17 as a potential therapeutic target to alleviate the burden of neuropsychiatric morbidities in infants surviving neonatal meningitis.

## Methods

### Bacterial strains and growth conditions

GBS BM110 strain (here referred to as GBS) is a bacterium belonging to the hyper-virulent clonal complex 17 (CC17), a well-known isolate from humans with invasive disease^81^. The isogenic non-haemolytic (βH/C-deficient) ΔcylE mutant (here referred to as CC17ΔcylE) was constructed by in-frame cylE gene depletion as previously described^82^. The GBS NEM316 strain (here referred to as non-CC17) is also a capsular serotype III strain, belonging to the CC23. Overnight (ON) cultures at 37□°C in Todd-Hewitt (TH) broth (Difco Laboratories) were 1:100 diluted in the same medium and bacteria were grown until mid-log phase (OD_600 nm_∼0.900), pellet and washed twice with phosphate-buffered saline (PBS). The OD_600nm_ was adjusted to ≈0.600±0.005, corresponding approximately to 2.0×10^8^ CFU/mL. Inocula were prepared by further dilution from this bacterial suspension. Inocula and organs were plated and ON grown in TH agar (Difco Laboratories) for bacterial count.

### Animals and ethics statement

C57BL/6 and *Tcrd^-^*^/-^ mice (originally obtained from The Jackson Laboratory) were bred and housed at the Instituto de Investigac□ão e Inovac□ão em Saúde (i3S) animal facility. *Il17a*^-/-^ mice were housed and bred at the Gulbenkian Institute for Molecular Medicine (GiMM Faculdade de Medicina da Universidade de Lisboa). The genetically modified mouse strains used are in the C57BL/6J genetic background and have been backcrossed at least ten times. All mice were maintained in specific pathogen-free conditions. Animal experiments complied with the ARRIVE guidelines, followed the recommendations of the European Convention for the Protection of Vertebrate Animals used for Experimental and Other Scientific Purposes (ETS 123) and Directive 2010/63/EU and Portuguese rules (DL 113/2013). All experimental protocols including animals were approved by the competent national authority Direcção Geral de Alimentação e Veterinária (DGAV), and by the respective Institute’s Animal Ethical Committee (ORBEA) (No 0421/2022-09-02). People directly working with animals are all certified by DGAV. All animals were kept in a controlled environment (20°C, 45–55% relative humidity) under a 12 h light/dark cycle, with free access to water and food. All efforts were made to minimise animal suffering and to reduce the number of animals used. Since our experiments were designed to colonise progenitors during pregnancy, we did not randomise their litters. No blinding was carried out.

### Mouse model of neonatal GBS meningitis

We adapted our intra-vaginal (i.vag.) mouse model of neonatal infection to the C57BL/6 mice^16^. Briefly, pregnant mice were i.vag. inoculated, using a micropipette, with 40 μL of GBS suspension containing 3×10^4^ cells, or with PBS, at the gestation days (G) 16 and 17. The presence of a vaginal plug was considered G1. Females were allowed to deliver spontaneously, and infected pups were kept with their mothers until weaning. C57BL/6 WT, *Tcrd*^-/-^ and *Il17a*^-/-^ were used. Both female and male pups were used unless otherwise stated.

### Survival, clinical scoring and organ colonisation

Pups born from i.vag. infected or control dams were monitored every 12 h, during the first indicated days of life for survival studies. A neonatal longitudinal clinical scoring, adapted from Fehlhaber *et al.*^83^, was used to evaluate the pup’s disease progression and death prediction during the first 5 days of life.

For the bacterial dissemination and assessment studies, the indicated tissues were collected in PBS, homogenised, serially diluted and plated on TH agar for CFU count.

### Flow cytometry

Animals were deeply anaesthetised by isoflurane inhalation, perfused with saline and tissues were aseptically removed. The meninges were collected in RPMI 1640 medium (HyClone GE Healthcare Life Sciences) with 2% FBS (heat-inactivated, Biowest) and 2.5 mg/ml collagenase D (Roche), and incubated at 37 °C for 30 min. The tissue is passed through a 1 mL syringe with a 25 G needle 5 times, washed twice with washing solution (RPMI 1640, 2% FBS, 2 mM EDTA), and passed through a 70 µm mesh filter. The meninges were centrifuged at 450 g, 5 min, at 4 °C, resuspended in RPMI 1640 with 2% FBS and counted. The brain was collected in RPMI 1640 medium, cut into small pieces (1–2 mm) and gently digested with 7.5 mg/mL collagenase D for 30 min. To aid dissociation, samples were pipetted up and down thoroughly every 10 min during the incubation period. Cells were then passed through a 100 μm cell strainer and washed. Stock isotonic Percoll (SIP, Cytiva) 90% was prepared with 10x HBSS without Ca^2+^ and Mg^2+^ (Gibco^TM^), and further diluted to 70%, 37% and 30% in 1x HBSS. Cell suspensions were suspended in 70% SIP. Each dilution of Percoll was layered in a 15 mL conical tube, and Percoll gradients were centrifuged at 800 g, 20 min, at 20 °C. The enriched population of leucocytes/microglia were carefully collected at the 70–30% interphase, washed and counted. The spleen and thymus were collected to RPMI 1640 with 2% FBS medium, gently dissociated in a 100 μm cell strainer to yield a single cell suspension, washed and counted.

A total of 5×10^5^ (meninges) or 1×10^6^ cells were plated in 96-well plates for staining. Suspensions were blocked with 5 μg/mL rat anti-mouse CD16/32 (eBiosciences) and 0.5 mg/mL normal rat serum for 10 min on ice prior to staining to reduce non-specific antibody binding. Cells were incubated with FVD eFluor 506 and then with the following fluorescence-coupled monoclonal antibodies, used at previously determined optimal dilutions: CD45 FITC (Clone 30-F11), CD69 PE (H1.2F3), CD8 PerCP-Cy5.5 (53-6.7), CD3 Pe-Cy7 (17A2), TCRδ APC (GL3), TCRβ eFluor 780 (H57-597, eBioscience), CD4 Pacific Blue (RM4-5), CD44 BV510 (IM7), Siglec-F PE (E50-2440, BD Biosciences), Ly6C PerCP-Cy5.5 (HK1.4), CD11b Pe-Cy7 (M1/70), Ly6G Pacific Blue (1A8), CD5 PE (53-7.3, BD Biosciences), TCRβ PE (H57-597, Invitrogen), CD19 APC-Cy7 (6D5), IFNγ PE (XMG1.2), TCRδ Pe-Cy7 (GL3), IL-17 APC (TC11-18H10.1), CD3 APC-Cy7 (17A2), Ly6G AF647 (1A8), Ki67 Alexa Fluor 488 (16A8), TCRVγ4 PE (UC3-10A6), CD45 PerCP-Cy5.5 (30-F11), TCRVγ1 APC (2.11), TCRβ eFluor 450 (H57-597, Invitrogen), CD62L PE (MEL-14), CD44 PerCP-Cy5.5 (IM7), PD-1 Alexa Fluor 647 (29F.1A12), CD69 BV421 (H1.2F3), CD4 BV421 (RM4-5), CD69 PE (H1.2F3), CD62L APC (MEL-14), TCRVγ4 PerCP-Cy5.5 (UC3-10A6), TCRVγ1 Pacific Blue (2.11), CD24 FITC (M1/69, Invitrogen), CD45RB Pacific Blue (C363-16A), CD69 Pe-Cy7 (H1.2F3), CD25 APC-Cy7 (PC61), Sirpα FITC (P84), CD200R PE (OX-110), MHC-II APC (M5/114.15.2), F4/80 APC-Cy7 (BM8) and/or CX3CR1 Pacific Blue (SA011F11). All antibodies are from Biolegend unless otherwise mentioned. For intracellular staining, cells were stimulated in complete RPMI 1640 with PMA (Phorbol 12-Myristate 13-Acetate; 50 ng/ml, Sigma-Aldrich) and Ionomycin (1 μg/mL, Sigma-Aldrich) and Brefeldin A (BFA) (10 μg/mL, Sigma-Aldrich), for 4 h at 37 °C, 5% CO_2_, before staining for extracellular markers. Cells were then permeabilised and stained using a Foxp3/Transcription Factor Staining Buffer Set (eBioscience). Cells were acquired in FACS Canto II (BD Biosciences) or Aurora (Cytek) flow cytometers. Post-acquisition analysis was performed using FlowJo Software v10 (Tree Star). For flow cytometry cell sorting, cells were stained with FVD eFluor 780 (eBioscience) and CD45 FITC (Clone 30-F11, Biolegend), and a minimum of 20,000 live CD45^+^ cells were sorted (FACS Aria II) into microcentrifuge tubes containing 0.04% BSA in DPBS, which were previously coated with 1 mL FBS, for 24 h. Samples were always kept on ice.

### Sample preparation for single-cell RNAseq (scRNAseq)

Male pups at P3 from infected and uninfected groups were deeply anaesthetised by isoflurane inhalation, perfused with ice-cold saline and the dural meninges were dissected and processed following adaptation from Scheyltjens and colleagues^86^. Briefly, the meninges were collected in RPMI and digested with 7.5 mg/mL of collagenase D for 35 min at 37 °C, with shaking at 300. Every 10 min, the samples were pipetted up and down thoroughly at room temperature. The meninges were then passed through a 100 µm mesh filter and pooled into 15 mL falcon tubes, separated by group. Cells were washed, transferred to a 1.5 mL polystyrene round-bottom tube with cap, and rinsed thrice with RPMI 1640. The samples were finally washed, resuspended in DPBS and counted. After cell sorting, cells were centrifuged and resuspended in DPBS with 0.04% BSA at a concentration of 1,000 cells / µL for chip loading. The GEM (Gel Bead-In Emulsion) barcode generation was performed according to the 10x Genomics instructions.

### scRNAseq analysis

Raw sequenced files were processed in Cell Ranger (v9.0.1) using the GRCm39-2024-A mouse reference. Seurat (v5.0.3) was used to import the Gene count matrices for each of the experiments. To clean the dataset we removed the ambient RNA counts using SoupX (v1.6.2) and cells with high mitochondrial gene percentage. Data was normalised and scaled following the Seurat default recommendation. Latent representations computed by scVI (v1.3) based on the top 2000 variable genes were used to build the integrated UMAP. Differentially expressed gene between clusters were identified Surat’s findallmarkers function. Gene signature scores were calculated using UCell (v2.4.0). For gene set enrichment analysis, deferentially expressed genes (DEGs) between clusters were identified using wilcoxauc function from presto. The GSEA and gseGO functions from clusterProfiler (v4.12.0) were used for the gene set enrichment analysis.

### scRNAseq analysis of IL-17R from adult mouse database

Previously published single-cell RNA sequencing data from the adult mouse brain^37^ were downloaded from http://dropviz.org and analysed using R. The dataset was analysed using the original pipeline described by Jin *et al*.^84^, including the default method for the computation of the communication probability and the use of 10 cells as the threshold for the minimum number of cells needed to express the receptor.

### Bulk RNA sequencing

Male pups at P3 were deeply anaesthetised by isoflurane inhalation, perfused with saline, and euthanised by decapitation. The meninges were collected and frozen at −80°C until RNA extraction. Two meningeal tissues were pooled before RNA isolation, and the pooled preparation constituted a single data point. Frozen tissues were retrieved on dry ice, and RNA was extracted using the Pure link^TM^ RNA Mini Kit (Ambion) with on-column DNase (Roche) treatment. Quantification and RNA integrity were assessed using an Agilent 2100 Bioanalyzer system. RNA integrity numbers (RIN) above 6.5 were used for library preparation and reverse transcription, followed by sequencing using Illumina PE150 technology (Novogene).

### Bulk RNA-seq data processing and analysis

RNA-seq reads were pre-processed in TrimGalore prior to alignment to remove low-quality and adapter sequences. Sequence alignment was carried in STAR v2.7.10a with the mouse GRCm39 reference using the Gencode vM33 gene models. Counts per gene were generated by STAR. A principal component analysis (PCA) was performed using the rlog-normalized count data to inform about the sample variability and structure across condition. Differential expression analysis was accessed in DESeq2 (v.1.34). Log2 fold change results were shrunk using the apeglm package to remove noise (DE genes with low counts and/or high dispersion values) while preserving significant differences. Gene set enrichment Pathway analysis was performed in ClusterProfiler (GSEA function) using a pre-ranked list of genes calculated using the signed log2 fold change times the -log10 of the p-value. Enrichment was calculated for the mouse hallmark gene sets that were retrieved from the Molecular Signatures Database (v2023.2.Mm).

### Cytokine analysis

Meningeal and splenic supernatants, upon single cell suspension isolation and 24 h stimulation with 50 ng/mL phorbol myristate acetate (PMA) and 500 ng/mL ionomycin, were collected and stored at −80 °C until use. The cytokines IL-6, IL-10 (R&D Systems), IL-17, IFN-γ and TNF-α, through ELISA, following Invitrogen or R&D Systems kits’ instructions, respectively.

Uninfected and Infected P3 splenic CD45^high^ cells were stimulated with GBS, at a multiplicity of infection (MOI) of 5 and 10, for 24 h. The cytokines IL-6, IL-10 (R&D Systems), IL-17 and TNF-α, through ELISA, following Invitrogen or R&D Systems kits’ instructions, respectively.

To study γδ T cell response to GBS, 2×10^4^ of splenic irradiated antigen-presenting cells (APCs) per well were incubated for 6 h with heat-killed GBS BM110 MOI 100 or RPMI.

Then, 5×10^3^ γδ T cell cells/well sorted from the lungs of P3 uninfected and infected pups were added and incubated for 72 h, with or without 10 ng/mL of recombinant mouse Il-1β and IL-23 (Biolegend). The supernatants were collected and stored at −80 °C until use. IL-1β, IL-23, IL-17 and IFN-γ were analysed through ELISA according to Invitrogen instructions.

### Intra-cisterna magna injections

P1 infected pups were wrapped in gauze and completely covered in ice for approximately 10 min. Newborns were injected via intra-cisterna magna with 1 mg/mL anti-mouse IL-17 or the IgG isotype control (InvivoMAb Biocell), diluted in 0.3 mg/mL of Fast Green, in a 2 µL of total volume per pup. The injection was performed using a 10 µL Hamilton syringe with a 30 G needle. The intra-cisterna magna was visually located through the thin clear skin, on the neck right below the skull. The needle was injected at 1.5 mm depth and the content was slowly injected. After injection, pups were allowed to completely recover on a heating pad under a red lamp before returning to the cage. CFU of meninges, brain, lungs and liver were analysed 24 h post-injection.

### Blood-brain barrier permeability assessment

For *in vivo* BBB permeability assessment, 0.3 mM of 3-5 KDa FITC-dextran (FD4, Sigma-Aldrich), in a 40 µL of total volume, were injected subcutaneously in P1 pups infected with GBS. Five minutes (min) after injection mice were deeply anesthetised with isoflurane. Blood was collected after the puncture of the right atrium and kept on ice. Blood was centrifuged at 10,000□×□g for 15 min at 4 °C and the serum was collected and stored at −80 °C until use. Mice were then perfused with PBS and the brain was extracted, divided into two hemibrains (hemispheres free of cerebellum and olfactory bulbs), weighted and immediately frozen on dry ice. The brains were stored at −80 °C until use. To measure the fluorescence intensity, the serum was diluted in PBS (20 µL of serum in 30 uL of PBS) and 100 uL of PBS was added to the hemibrain to homogenise the tissue. The fluorescent intensity was measured using the SpectraMax iD3 (Molecular Devices). The degree of BBB disruption was described as the permeability index^85^.

### Developmental milestones

Pups born from uninfected and Infected C57Bl/6 WT and *Tcrd^-/-^*mice were assessed daily for weight since P0 and the acquisition of developmental milestones since P1, as previously described^86^. Pups were gently removed from their cages and placed on a heating pad and under a red lamp, while the litter was tested. Pups were weighted daily and visually examined for eye-opening. Each day the pups were evaluated through a battery of tests assessing the attainment of milestones appropriate for their age^86^. After testing, the pups returned to their cages. The development milestones are as follows:

#### Surface righting (P1-P13)

Pups were individually held on their backs (supine position), with paws facing upwards. The time pups took to flip over onto their abdomens (prone position) was noted. The test was terminated if the pup failed to turn over within 30 seconds or when it could turn itself in less than 1 second for two consecutive days.

#### Negative geotaxis (P1-P14)

Pups were individually placed on a wire mesh set at a 45° angle with heads facing downwards. The time to turn around by 180° and move upwards was recorded. The test was repeated daily until pups passed in < 30 seconds for two consecutive days.

#### Cliff aversion (P1-P14)

Pups were positioned on the edge of a pipette tip’s box with forepaws and the snout hanging over the edge. The time taken to turn and crawl away from the edge was noted. The test was repeated daily until pups passed in < 30 seconds for two consecutive days.

#### Rooting (P1-P12)

Pups were gently rubbed with the smooth tip of a brush along one side of the head. A positive response was noted if the pup moved its head toward the filament. The test was repeated daily until the pups responded correctly for two consecutive days.

#### Forelimb grasp (P4-P14)

Pups were held with their forelimbs grasping a wire bar, while their hindlimbs were not supported on any surface. The time pups were able to remain suspended was noted. The test was repeated daily until pups could stay suspended for a minimum of 1 s for two consecutive days.

#### Auditory startle (P7-P18)

Pups were individually exposed to a hand clapping, at approximately 25 cm distance. The day the pups responded with a quick involuntary jump was noted.

#### Ear twitch (P7-P15)

The fine filaments of a brush were gently touched against the tip of the pup’s ear. Positive responses were noted if pups responded by flattening their ears. The test was repeated daily until the pups responded correctly for two consecutive days.

#### Open field traversal (P8-P21)

Each pup was placed at the centre of a circular 13 cm diameter arena. The latency to move out of the arena was recorded. The test was repeated daily until pups walked out of the arena in < 30 seconds for two consecutive days.

#### Air righting (P8-P21)

Pups were held on their backs (with four paws turning upwards), 10 cm above a cotton pad, and released. The test was repeated daily until pups landed on all four paws for two consecutive days.

### Behavioural tests

Before the performance of the behavioural tests, P42 control or survivor mice were allowed a 2-week adaptation period to the behavioural test facilities. Tests were conducted in the dark (active) cycle. The order of the tests was as follows: elevated plus maze, social test, open field and novel-object recognition. The social test was performed approximately 4 h after the elevated plus maze. The remaining tests were performed with an interval of 24 h. All materials were cleaned with a neutral detergent without smell between animals, to prevent olfactory cues from influencing behavioural responses. The same groups of animals performed all tests (a 3R strategy). All tests were videotaped with a camera positioned above the apparatus. Student’s t-test was used to assess genotype and group effects, and the animals were separated by sex.

#### Elevated plus maze (EPM)

EPM apparatus was placed 50□cm above the ground and is composed of a cross form with two open arms (30 × 5 cm) and two closed arms (30 × 5 cm, surrounded by 15□cm-high opaque walls), and was used to evaluate anxiety levels. Identical arms are in opposite positions to each other, with the arms emerging from a central platform (5 × 5 cm). The test was initiated by placing the mouse in the centre of the apparatus facing an open arm and allowing it to move freely for 5□min. The lights were on during the test. The analyses were performed using a tracking system (Smart Video Tracking Software v 2.5, Panlab).

#### Social test

The apparatus, used for sociability preference studies, consisted of a wide acrylic box divided into three compartments (two 33 × 33 cm side compartments and one 12 × 12 cm central compartment) connected by open doors. The social tests consisted of three phases: the habituation phase, where the animal is positioned in the central compartment and allowed to explore for 5 min; the target phase, where one side compartment has an empty cage (round and gridded) and the other side compartment has the same cage with an animal inside (of the same age and sex as the sample animal), and the animal is positioned in the central one and allowed to explore for 10 min; the novel vs familiar phase, where an animal is added to the empty cage and the sample animal is positioned in the central compartment and allowed to explore for another 10 min. Animal/cage exploration was defined as the mouse nose touching or directed towards the object at a distance shorter than 2□cm. The analyses were performed in the Observer 7 XT software (Noldus Information Technology, Wageningen, The Netherlands).

#### Open field

The open field test was performed to evaluate activity levels, anxiety, as well as the apparatus habituation period for the novel object recognition test. Each mouse was positioned in the centre of an opaque arena (43 × 43 cm) and allowed to move freely for 10□min. The total distance travelled, peripheral activity (locomotion along the walls) and centre activity (locomotion in the central zone) were automatically obtained through video tracking (Smart Video Tracking Software v 2.5, Panlab).

#### Novel object recognition

The NOR test was used to evaluate memory/cognition. It was performed in an arena (43 × 43 cm), consisting on three phases: the habituation phase, in which mice were allowed to explore the apparatus for 10□min; the acquisition/sample phase 24□h after the habituation, where mice were placed in the apparatus with two identical objects (familiar object) for 10□min; and the retention/choice session, performed 4□h later, in which a novel object and a familiar object were used, and mice were allowed to explore the objects for 3□min. The objects chosen for this experiment were approximately the same height and weight, with different shapes, colours or materials. Object exploration was defined as mice nose touching or directed towards the object at a distance shorter than 2□cm, and the duration of time mice spent exploring each object was recorded by the observer, using Observer 7 XT software (Noldus Information Technology, Wageningen, The Netherlands). The recognition index was calculated as the ratio T_N/F_ / (T_N_ + T_F_) [T_N_ = time exploring the novel (N) object; T_F_ = time exploring the familiar (F) object]. As exclusion criteria, mice exploring the objects for periods lower than 10 s at the 3 min choice session were excluded from the analysis.

### Brain Tissue Processing and Immunostaining

For immunohistochemistry analysis, animals were deeply anaesthetised and perfused with PBS. The brains were removed, post-fixed by immersion in 4% PFA for 24□h, at 4□°C, washed with PBS and then cryoprotected using sucrose 30%. Samples were thereafter washed in PBS and mounted in OCT (Thermo Scientific) embedding medium, frozen and cryosectioned in the CM3050S Cryostat (Leica Biosystems). Brain coronal sections (30 μm) were collected by free-floating, in PBS, transferred to cryoprotectant solution (30% sucrose and 30% ethylene glycol in 0.1 M phosphate buffer) and stored at −20□°C until staining. For immunolabeling, brain slides were permeabilised with 0.25% Triton for 10 min, blocked for 1 h in PBS with 0.1% Triton and 5% Bovine Serum Albumin (BSA) and incubated overnight (4 °C) with the primary antibodies: anti-Iba-1 (FUJIFILM Wako; 1:500), anti-NeuN (Sigma-Aldrich; 1:500) and anti-CD68 (Bio-Rad; 1:500). The following secondary antibodies were used: goat anti-rat IgG F(ab’)2 Alexa Fluor 488 (Invitrogen; 1:1000), goat anti-rabbit Alexa Fluor 568 (Invitrogen; 1:1000) and goat anti-mouse Alexa Fluor 647 (Invitrogen; 1:1000) for 1h30 at RT. DAPI (Thermo Fisher Scientific; 1:100) was used to mark the nuclei. Sections were mounted using a Fluorescence mounting medium (Dako), sealed with nail polish and stored at −20 °C.

### Primary Microglia Cell Cultures and Immunohistochemistry

Primary mixed glial cultures were obtained from newborn Wistar Han rat pups at P1 – P2. The pups’ brains were dissected in Hank’s Balanced Salt Solution (HBSS, Gibco^TM^) with 1% Penicillin-Streptomycin (P/S). After complete tissue dissociation, enzymatic digestion was completed using DNAse I (0.1 U/mL) (Irvine) and 0.25% trypsin-EDTA (Gibco^TM^). The cells were suspended in Dulbecco’s Modified Eagle Medium (DMEM) (1x)/Glutamax (Gibco^TM^) supplemented with 10% of FBS and 1% of P/S. Cells were seeded in T75 flasks coated with 0.01 mg/mL of PDL at a density of two brains/flask. The cell cultures were maintained for 10 days at 37 °C in a humidified incubator with 5% CO_2_. The medium was partially replaced at day 4 and then totally replaced every 2 days. MGCs were shaken at 200 rpm for 2 h at 37 °C to obtain purified microglia cells. The cell suspension obtained was centrifuged at 1200 rpm for 10 min and resuspended in DMEM/ Nutrient Mixture F-12 (DMEM/F-12) (GibcoTM) supplemented with 10% of FBS and 1% of P/S. Purified microglia were plated at a density of 0.5 x 10^5^ / cm in a 24-well plate with rounded coverslips in the wells.

For stimulation assays, microglia were cultured in medium alone or in the presence of heat-killed GBS at a multiplicity of infection (MOI) of 5. Each condition was set in duplicate, and cultures were maintained for 24 h at 37 °C with 5% CO_2_. Cells in coverslips were fixed with 4% PFA (Sigma-Aldrich) for 10 min at RT. Then, cells were permeabilised with 0.2% Triton X-100 (Sigma-Aldrich) for 10 min at RT and blocked with 3% BSA (Sigma-Aldrich) for 1h at RT. Fixed cells were incubated with primary antibody anti-iNOS (1:200, Abcam) overnight at 4 °C. Cells were washed and incubated with secondary antibodies (1:1000, Invitrogen) for 1h at RT, and nuclei stained with 1 μg/mL Hoechst 33342 (Sigma-Aldrich), for 10 min at RT. Coverslips were mounted with fluorescent mounting media (Dako).

### Imaging acquisition and quantification

Images were acquired in a Leica TCS SP8 confocal microscope (Leica Microsystems), using a PL APO 63x /1.30 Glycerol immersion objective for the Iba-1 and CD68 quantification, and a PL APO 10x /0.40 CS2 objective for NeuN quantification. The prefrontal cortex, the motor cortex and the hippocampus were studied. A total of 3 slices per brain were acquired. In each slice, 5 images were taken in the cortex using each objective, and 6 and 12 images of hippocampus regions (CA1, CA2, CA3 and Dentate Gyrus) were acquired with the 10x and the 63x objective, respectively. For the analysis, Fiji software was used for microglia and neuron quantification. The quantification of neurons was automatically done with a custom-made ImageJ/Fiji script from the Advanced Light Microscopy platform at i3S. Three-dimensional microglia reconstruction was performed using Imaris software (version 9.1.1, Bitplane). For microglia cultures, imaging was performed using a Zeiss AxioImager Z1 fluorescence microscope equipped with an Axiocam MR v3.0 camera, an HXP 120 light source, and a 40x/1.30 objective. The fluorescence intensity of microglial cells was quantified using ImageJ software.

### Statistical analysis

All data were analysed with the GraphPad Prism software (v.9, GraphPad software Inc. CA). Means and standard errors of the means (SEM) were calculated and correspond to the indicated independent experiments. The log-rank (Mantel–Cox) test was used to analyse the survival curve. Differences between multiple groups were analysed by one-way or two-way ANOVA, whenever appropriate, with Sidak’s Multiple Comparison Test, for α□=□0.05. For microglia morphology analysis by immunocytochemistry or for cell cultures analysis, a linear mixed model analysis was performed using SAS® Visual Statistics with REML function, followed by unpaired Student’s t test, using the animal or the culture as a random variable, respectively. All other comparisons were done with an unpaired Student’s t-test or one-way ANOVA, as indicated in the figures. Normality was verified by the Shapiro–Wilk normality test. Homogeneity of variance was estimated by an F test. Assumptions of sphericity in ANOVA were also verified. The minimum sample size was determined using G*Power analysis for a two-tailed t-test comparing the mean and standard deviation of two independent groups with effect size d□=□7, α□=□0.05 and power of 0.95. CFU data were log10 transformed. The number of biological replicates (n) and the number of independent experiments are indicated in the figure legends. Significance was represented by the following symbols: *P□<□0.05, **P□<□0.01, ***P□<□0.001, ****P < 0.0001.

## Acknowledgements

The authors acknowledge the support of the personnel in the animal facilities and the i3S scientific platforms Advanced Light Microscopy, a member of the national infrastructure PPBI-Portuguese Platform of BioImaging (supported by POCI-01-0145-FEDER-022122), and Translational Cytometry (TraCy). The authors also thank Ricardo Vieira for his assistance with published single-cell RNA sequencing data analyses.

This work was supported by National Funds through Fundac□ão para a Cie□ncia e a Tecnologia (FCT), I.P., under the project EXPL/SAU-INF/1217/2021. I.L. is supported by FCT through a PhD fellowship (2020.05394.BD). E.B.A. is funded by FCT through *Estímulo Individual ao Emprego Científico* (CEECIND/03675/2018). A.H. is supported by the Hannover Biomedical Research School (HBRS), the Center for Infection Biology (ZIB) and by the Deutsche Forschungsgemeinschaft (DFG, German Research Foundation) grant number GE3062/2-1 to H.G.

## Author Contributions

Conceptualisation, E.B.A.; Methodology, I.L., J.B., A.M., J.R. and E.B.A.; Investigation, I.L., J.S., J.B., P.M., N.R., J.R., A.H., M.M., B.C.; Formal Analysis, I.L., J.B., A.H., B.C., A.M., M.V., J.R. and E.B.A.; writing - original draft preparation, I.L. and A.H.; writing - review and editing, T.S., H.G., M.V., J.R. and E.B.A.; Supervision, E.B.A.; Funding acquisition and resources, E.B.A.

---

**Figure S1. Immune kinetic analysis in the meninges and spleen of GBS-infected newborn mice.** Pregnant C57BL/6 mice were intravaginally colonised with 3×10^4^ CFU of GBS hypervirulent strain BM110, at gestational days 16 and 17. Control animals received PBS. Newborn mice were sacrificed at indicated postnatal days (P). **(a-f)** Frequency and number of indicated populations in the meninges. Each symbol represents a single pup. Error bars indicate the mean ± SEM (n = 8-12 Control, n = 5-10 Meningitis). Comparisons by Student’s t-test within each time point. **(g-l)** Frequency and number of indicated populations in the spleen. Each symbol represents a single pup. Error bars indicate the mean ± SEM (n = 8-16 Control, n = 7-10 Meningitis). Comparisons by Student’s t-test within each time point. Statistical differences (P values) between groups are indicated; *P < 0.05, **P < 0.01, ***P < 0.001, ****P < 0.0001.

**Figure S2. Immune kinetic analysis in the brain of GBS-infected newborn mice.** Pregnant C57BL/6 mice were intravaginally colonised with 3×10^4^ CFU of GBS hypervirulent strain BM110, at gestational days 16 and 17. Control animals received PBS. Newborn mice were sacrificed at indicated postnatal days (P). **(a-f)** Frequency and number of indicated populations. Each symbol represents a single pup. Error bars indicate the mean ± SEM (n = 4-12 Control, n = 4-5 Meningitis). Comparisons by Student’s t-test within each time point. Statistical differences (P values) between groups are indicated; *P < 0.05, **P < 0.01, ***P < 0.001, ****P < 0.0001.

**Figure S3. Meningeal cytokines and** γδ **T cells systemic characterisation in neonatal meningitis.** Pregnant C57BL/6 mice were intravaginally colonised with 3×10^4^ CFU of GBS hypervirulent strain BM110, at gestational days 16 and 17. Control animals received PBS. Newborn mice were sacrificed at postnatal (P) day 3. **(a-d)** Characterisation of splenic γδ T cells. **(a)** Frequency of γδ T cells in the spleen of male and female pups at P3. Each symbol represents one pup (n = 8 Control, n = 4-5 Meningitis). Error bars indicate mean ± SEM. Comparisons by Student’s t-test. **(b)** Bee swarm plots for KNN smoothed UCell signature enrichment score of genes in https://www.wikipathways.org/pathways/WP5242. **(c)** Supernatant concentrations of indicated cytokines from *ex vivo* cultured meningeal cells, isolated from P3 pups, after 24 h of PMA and ionomycin stimulation. Each symbol represents a pool of 2 pups (n = 6-12). Data are presented as mean ± SEM. Comparisons by Student’s t-test within PMA/ionomycin absence or presence. **(d)** Frequency of IL-17 (top panel) and IFN-γ (bottom panel) produced by splenic cells at P3. Each symbol represents one pup (n = 6 Control, n = 4 Meningitis). Error bars indicate mean ± SEM. Comparisons by Student’s t-test between groups. **(e)** Supernatant concentrations of indicated cytokines from *ex vivo* cultured splenic cells, isolated from P3 pups, after 24 h of PMA and ionomycin stimulation. Each symbol represents a pool of 2 pups (n = 6-12). Data are presented as mean ± SEM. Comparisons by Student’s t-test within PMA/ionomycin absence or presence. **(f)** Frequency of Ki67 in meningeal γδ T cells. Each symbol represents one pup (n = 7 Control, n = 8 Meningitis). Error bars indicate mean ± SEM. Comparisons by Student’s t-test. **(g-i)** MFI and frequency of indicated surface markers in meningeal γδ T cells. Each symbol represents one pup (n = 3). Error bars indicate mean ± SEM. Comparisons by Student’s t-test. **(i)** Representative gating strategy (left panel) and frequency of indicated populations (right panel). Each symbol represents one pup (n = 3). Error bars indicate mean ± SEM. Comparisons by Student’s t-test. **(j)** Supernatant concentrations of indicated cytokines from *ex vivo* culture pulmonary γδ T cells, isolated from P3 pups, after 72 h of stimulation with GBS in the presence or absence of IL-1β and IL-23 and/or antigen-presenting cells (APC, splenic CD45^+^ irradiated cells). Each symbol represents a pool of 2 pups (n = 3-5 pools). Data are presented as mean ± SEM. Comparisons by One-way ANOVA. **(k)** IL-1β and Il-23 concentrations from *ex vivo* culture of splenic APCs upon 72 h GBS stimulation. Each symbol represents a pool of 2 pups (n = 3 pools). Data are presented as mean ± SEM. Comparisons by Student’s t-test. Statistical differences (P values) between groups are indicated; *P < 0.05, **P < 0.01, ***P < 0.001. DL, detection limit.

**Figure S4. Lung** γδ **T cell subsets analysis.** Pregnant C57BL/6 mice were intravaginally colonised with 3×10^4^ CFU of GBS hypervirulent strain BM110, at gestational days 16 and 17. Control animals received PBS. Frequency (left) and number (right) of lung Vγ1^+^, Vγ4^+^ and Vγ6 (Vγ1^-^Vγ4^-^) γδ T cells, at postnatal day (P) 3. Each symbol represents one pup (n = 4). Error bars indicate mean ± SEM. Comparisons by Student’s t-test within subsets.

**Figure S5. GBS BM110 brain load.** Pregnant C57BL/6 mice were intravaginally colonised with 3×10^4^ CFU of GBS hypervirulent strain BM110, at gestational days 16 and 17. Control animals received PBS. Bacterial load in the brain of 2 week-(2 wk) and 6-week-old (6 wk) survivors. Each symbol represents a single animal (n = 9 2 wk, n = 3 6 wk). Data is presented as mean. DL, detection limit.

**Figure S6. Immune composition of the meninges and spleen of GBS-infected newborn *Tcrd^-/-^* mice.** Pregnant *Tcrd*^-/-^ mice were intravaginally colonised with 3×10^4^ CFU of GBS hypervirulent strain BM110, at gestational days 16 and 17. Control animals received PBS. Newborn mice were sacrificed at postnatal day (P) 3. **(a-f)** Frequency and number of indicated populations in the meninges. Each symbol represents a single pup. Error bars indicate the mean ± SEM (n = 6 Control, n = 4 Meningitis). Comparisons by Student’s t-test. **(g-l)** Frequency and number of indicated populations in the spleen. Each symbol represents a single pup. Error bars indicate the mean ± SEM (n = 6 Control, n = 4 Meningitis). Comparisons by Student’s t-test.

**Figure S7. Expression of IL-17 receptor in different brain regions. (a-g)** Predicted expression of the IL-17 receptor chains a (*Il17ra*) and c (*Il17rc*) in the different cell types, in the indicated brain regions, using previously published single-cell RNA-seq data. The size of each dot represent the average expression of cells expressing each gene.

**Figure S8. Female survivors of neonatal GBS meningitis do not present long-term sequelae.** Pregnant C57BL/6 WT or *Tcrd^-/-^* mice were intravaginally colonised with 3×10^4^ CFU of GBS hypervirulent strain BM110, at gestational days 16 and 17. Control animals received PBS. Females born from the indicated groups were allowed to reach adulthood and submitted to behavioural tests. **(a-e)** Behavioural analysis in the open-field test. **(a)** Representative scheme of the open field test. The total distance covered **(b)**, the average speed **(c)** and the percentage **(d)** and time **(e)** of distance covered by zone were used as a measure of hyperactivity and anxious-like behaviour. **(f-i)** Cognitive test through novel object recognition (NOR). **(f)** Representative scheme of the novel object recognition test. The recognition index **(g)** was used as a measure of cognitive function. The object exploration time **(h)** was used as a control of the test. **(i)** Object exploration time in the familiar (F) and novel (N) object. **(j-m)** Elevated Plus Maze test. **(j)** Representative scheme of the elevated plus maze test. The total distance covered **(k)**, and the percentage of distance **(l)** and time **(m)** covered by zone were used as a measure of anxious-like behaviour. **(n)** The socialisation index in motivation and preference phases **(i)** was used as a measure of social preference. Each symbol represents a single animal (n = 9 Control, n = 9-10 Survivor). Data represented as mean. Comparisons by Student’s t-test within genotypes and social conditions.

**Figure S9. Behavioural assessement in male survivors of neonatal GBS meningitis.** Pregnant C57BL/6 WT or *Tcrd^-/-^*mice were intravaginally colonised with 3×10^4^ CFU of GBS hypervirulent strain BM110, at gestational days 16 and 17. Control animals received PBS. Male animals born from the indicated groups were allowed to reach adulthood and submitted to behavioural tests. **(a-d)** Behavioural analysis in the elevated plus maze test. **(a)** Representative scheme of the Elevated Plus Maze test. The total distance covered **(b)**, and the percentage of distance **(c)** and time **(d)** covered by zone were used as a measure of anxious-like behaviour. **(e)** The percentage of time covered in the indicated regions of the open field test. **(f-g)** Social test.**(f)** Representative scheme of the social test. The socialisation index in motivation and preference phases **(g)** was used as a measure of social preference. **(h-k)** Cognitive test through novel object recognition (NOR). **(h)** Representative scheme of the novel object recognition test. The recognition index **(i)** was used as a measure of cognitive function. **(j)** The object exploration time was used as a control of the novel-object recognition test. **(k)** Object exploration time in the familiar (F) and novel (N) object. Each symbol represents a single animal (n = 10 Control, n = 7 Survivors). Error bars indicate mean ± SEM. Comparisons by Student’s t test within genotypes and social conditions. Statistical differences (P values) between groups are indicated. *P < 0.05, **P < 0.01, ***P < 0.001.

**Figure S10. Meningeal RNA-seq analysis reveals altered cellular development in neonatal meningitis.** Pregnant C57BL/6 mice were intravaginally inoculated with 3×10^4^ CFU of GBS hypervirulent strain BM110 at gestational days 16 and 17. Control animals received PBS. **(a)** The meninges were isolated from the brain of male pups with meningitis and controls at postnatal day (P) 3, and total RNA was extracted and sequenced. **(b)** PCA plot showing segregation between uninfected controls (grey) and neonatal meningitis (yellow) meningeal transcriptomes. **(c)** Volcano plot depicting the gene expression differences in the meninges of pups with meningitis compared to uninfected controls. Red dots represent genes expressed at higher levels in neonatal meningitis, while blue dots represent genes with higher expression levels in healthy controls. Grey dots represent genes bellow the cutoff of significant (|log2 fold change| ≥ 0.5 and adjusted P values ≤ 0.05). Differential expression was evaluated using the Wald’s test, followed by adjustment for multiple testing with the Benjamini-Hochberg correction. Log2-fold change values were shrunk with the apeglm method to increase the signal-over-noise ratio of the effect size. **(d-e)** Dot plot of gene set enrichment analysis obtained by functional enrichment of differentially expressed genes in the meninges during neonatal meningitis. The diameter of the dot indicates the degree of significance of the ontology term. Red dots represent terms enriched in pups with meningitis, while blue dots represent terms enriched in healthy mice. **(f)** Heat maps depicting the relative expression values (z-score of each gene across samples) of the top seven gene pathways associated with cellular development. Only differentially expressed genes were represented, excluding genes belonging to more than one pathway. Individual RNA samples result from meninges pooled from 2 pups over 3 independent experiments (n = 3 pools per group).

**Figure S11. Microglia and neuronal quantification in male survivors of neonatal GBS meningitis.** Pregnant C57BL/6 mice were intravaginally inoculated with 3×10^4^ CFU of GBS hypervirulent strain BM110 at gestational days 16 and 17. Control animals received PBS. The male progeny was sacrificed 6 weeks after birth. **(a)** Representative images of NeuN and Iba1, and Imaris 3D reconstruction of microglia and CD68 in the cortex of 6-week-old animals from both groups. Scale bar 20 μm. **(b)** Quantification of NeuN cells in the brain’s somatosensory cortex (St cortex) and hippocampus. Comparisons by Student’s t-test. **(c)** Quantification of Iba-1 cells in the brain’s somatosensory cortex (St cortex) and hippocampus. Each symbol represents one animal (n = 4). Error bars indicate mean ± SEM. Comparisons by Student’s t-test. Microglia surface area **(d)** and volume **(e)** at the indicated brain regions. Each coloured symbol represents one animal (n = 4) and each background grey symbol represents one cell (n = 41-82 Control, n = 46-48 Survivor). Comparison by Nested t-test.

