## Supplemental figures for "Meningeal γδ T cells facilitate bacterial entry into the brain in neonatal meningitis and trigger long-term behavioural sequelae"

### Suplemmentary Figure 1

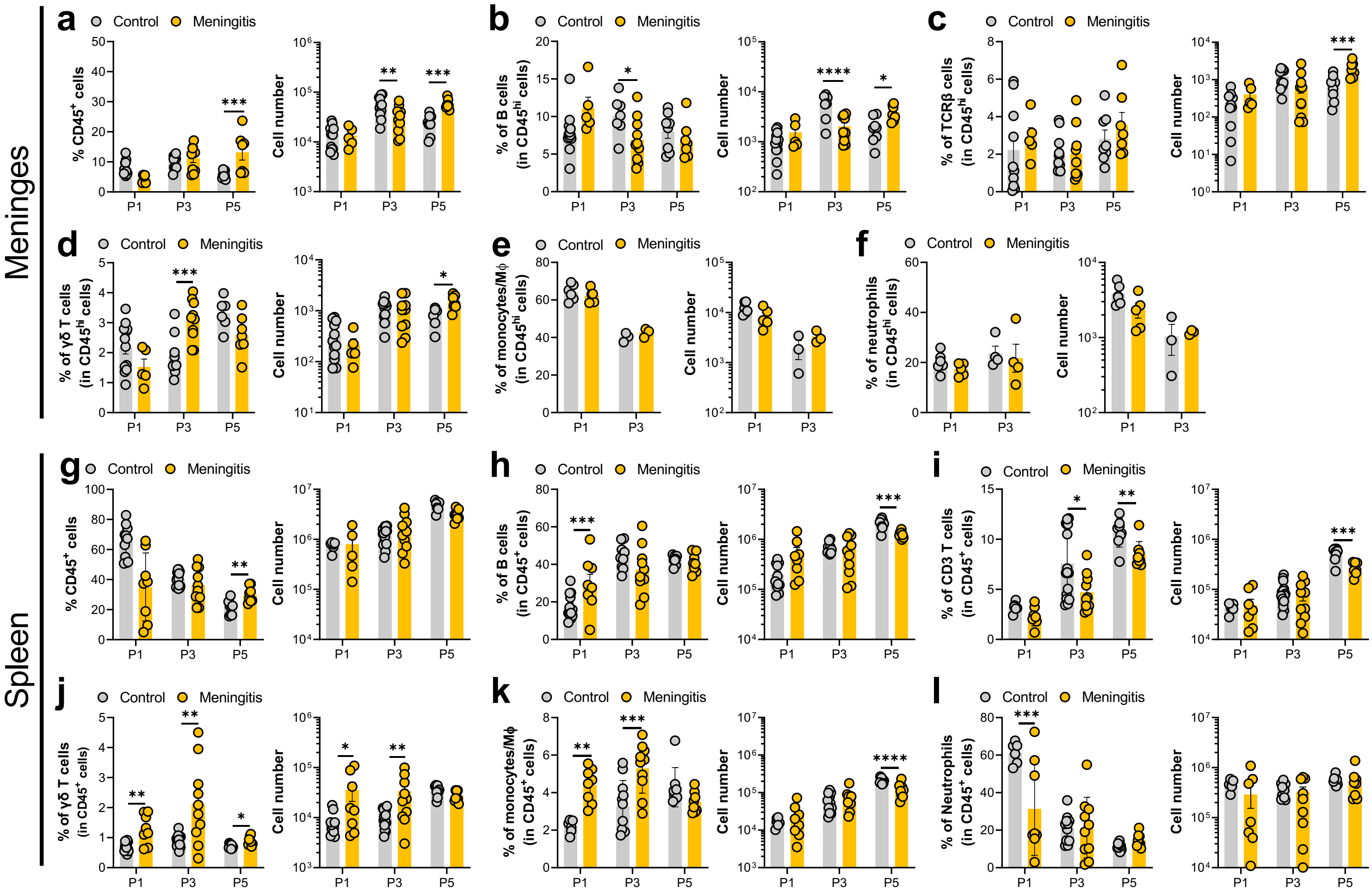

### Suplemmentary Figure 2

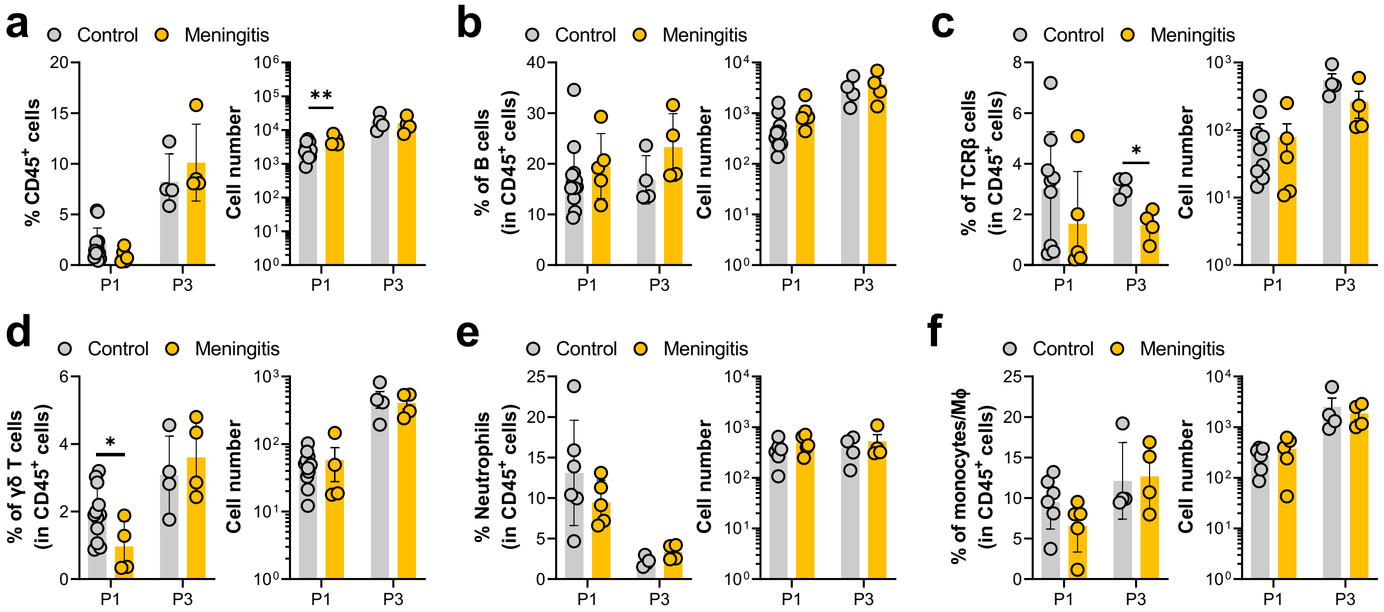

### Suplemmentary Figure 3

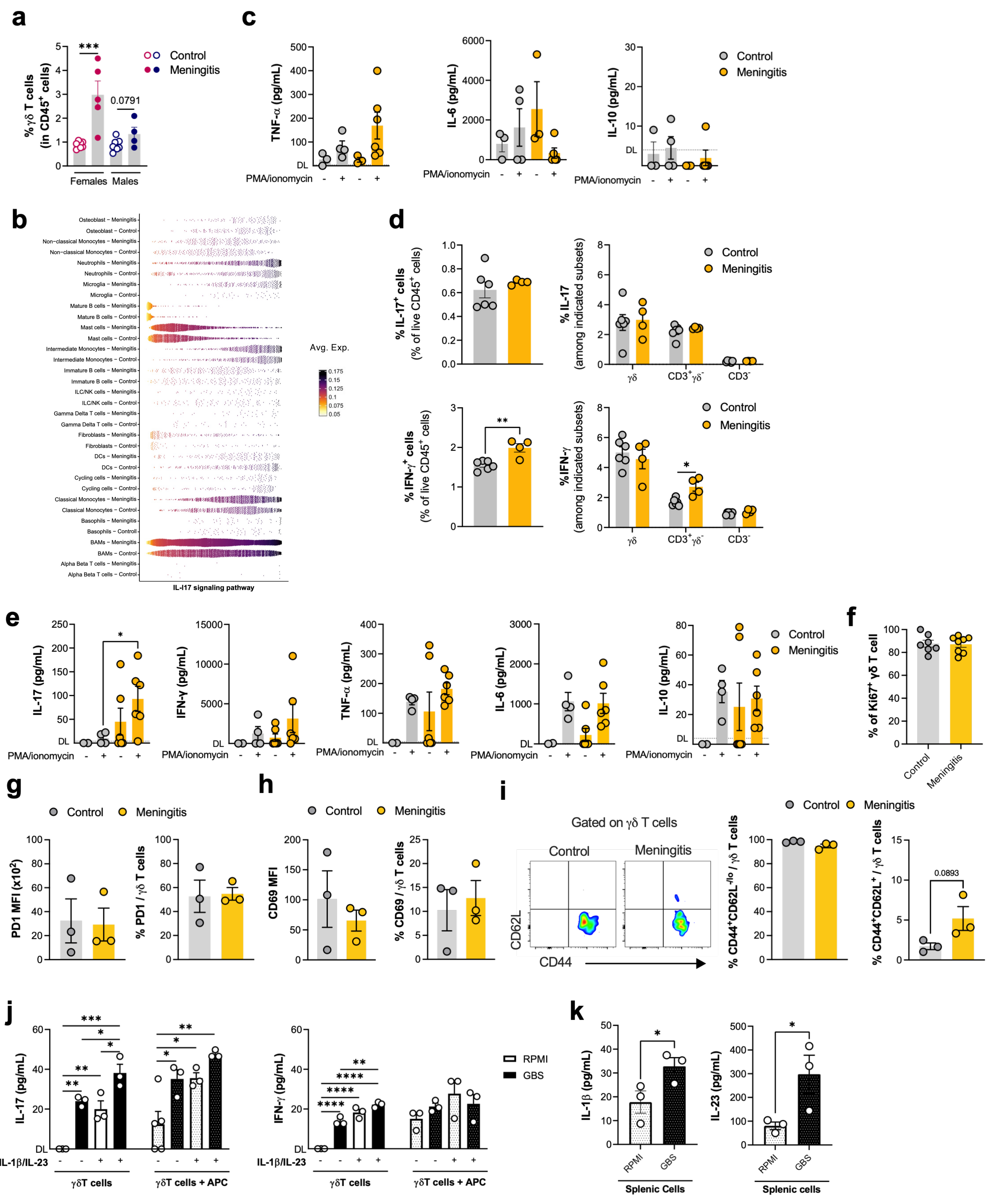

### Suplemmentary Figure 4

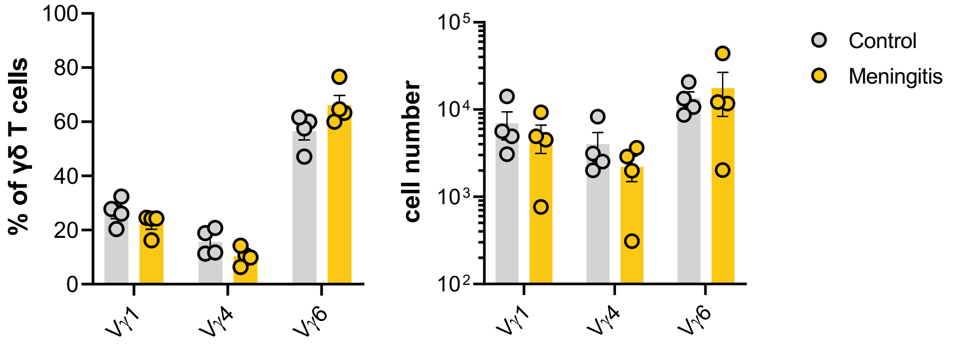

### Suplemmentary Figure 5

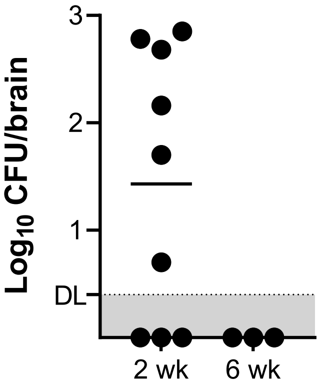

### Suplemmentary Figure 6

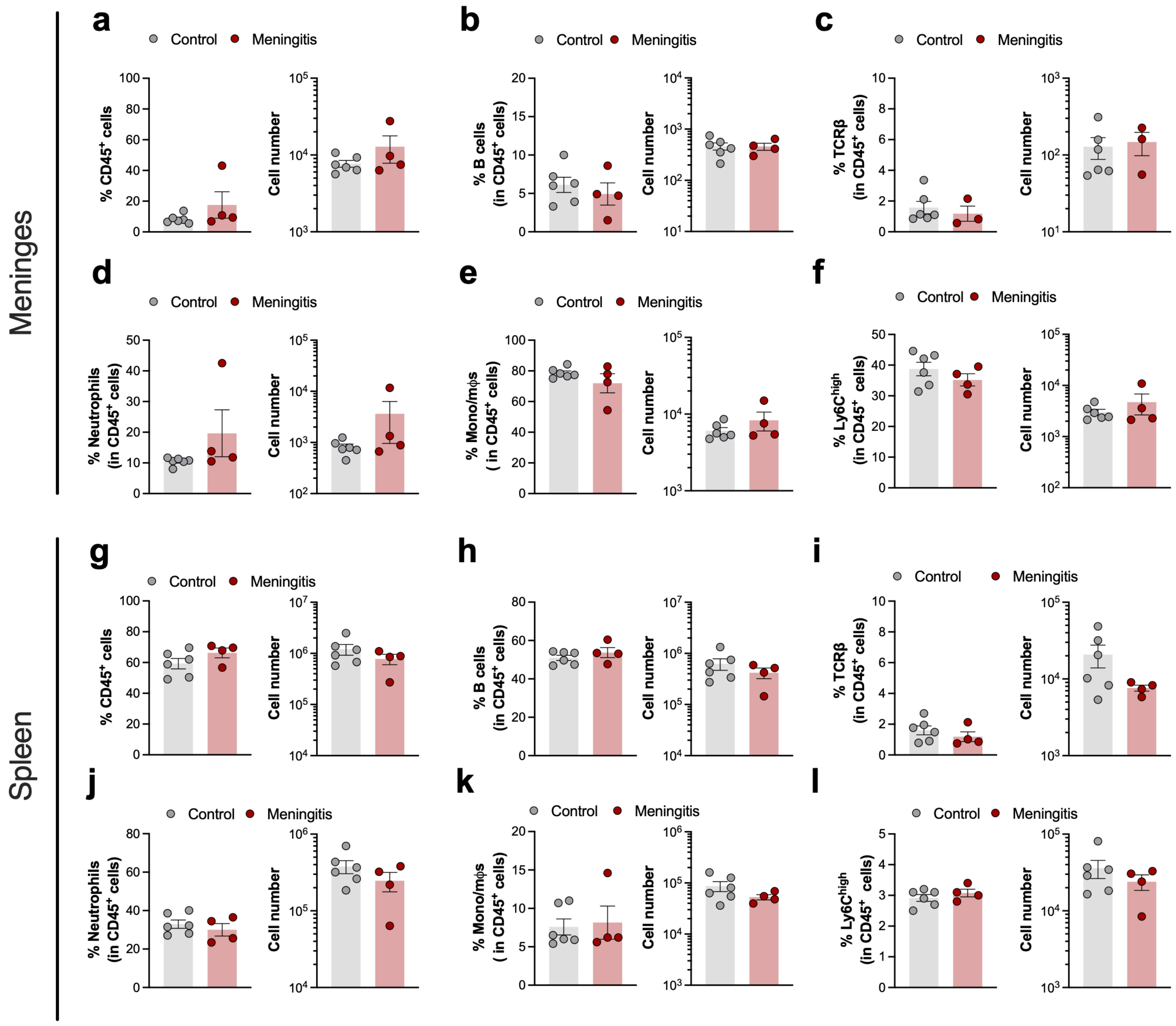

### Suplemmentary Figure 7

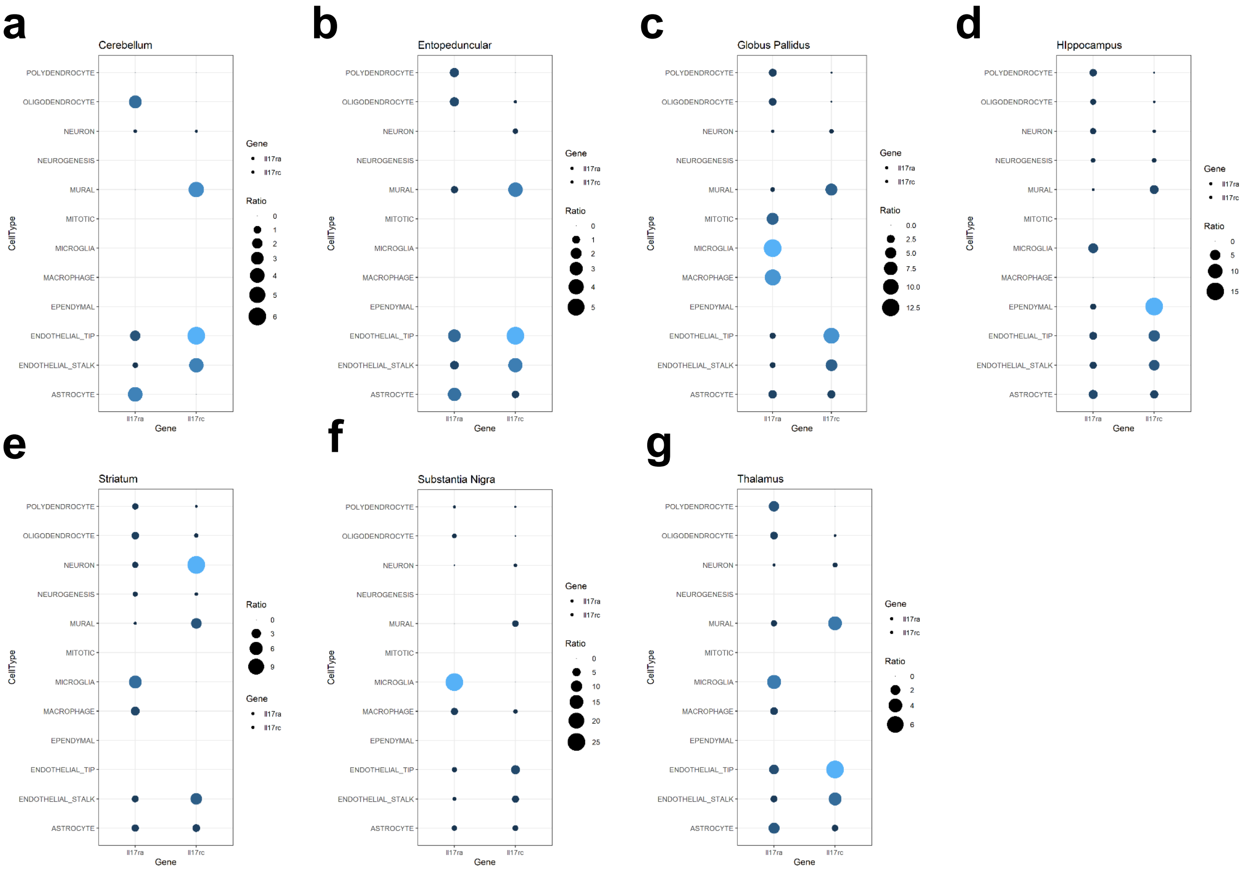

### Suplemmentary Figure 8

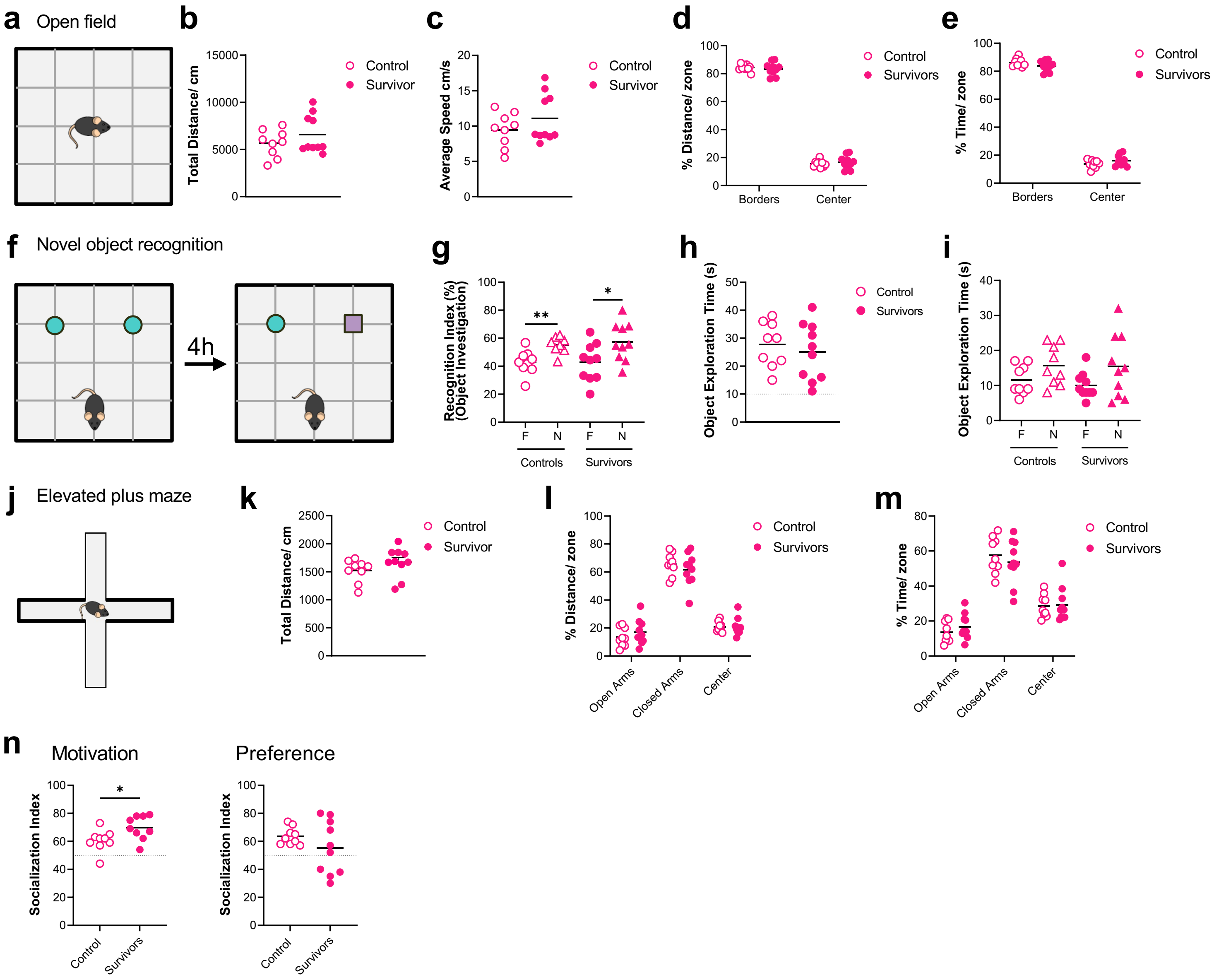

### Suplemmentary Figure 9

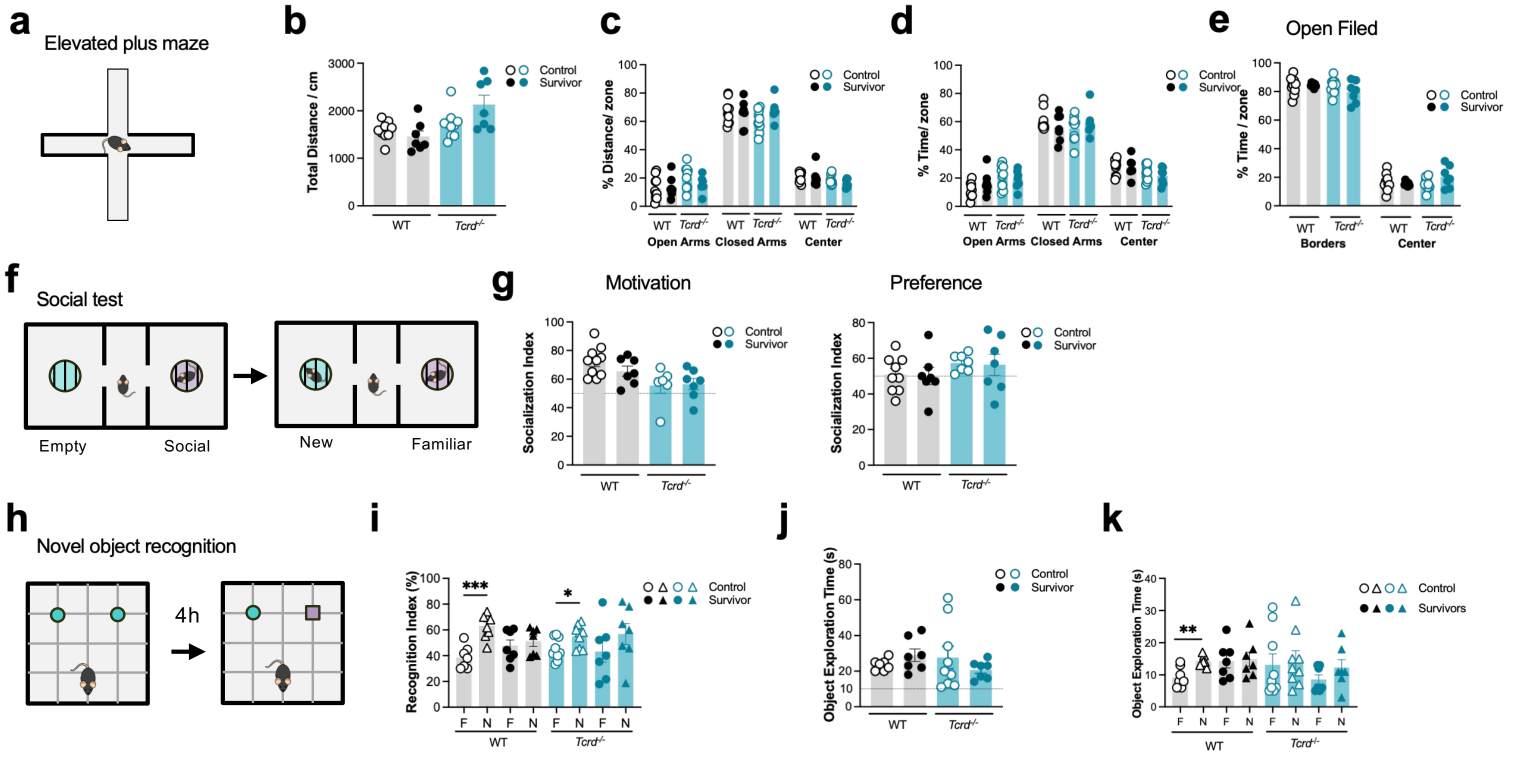

### Suplemmentary Figure 10

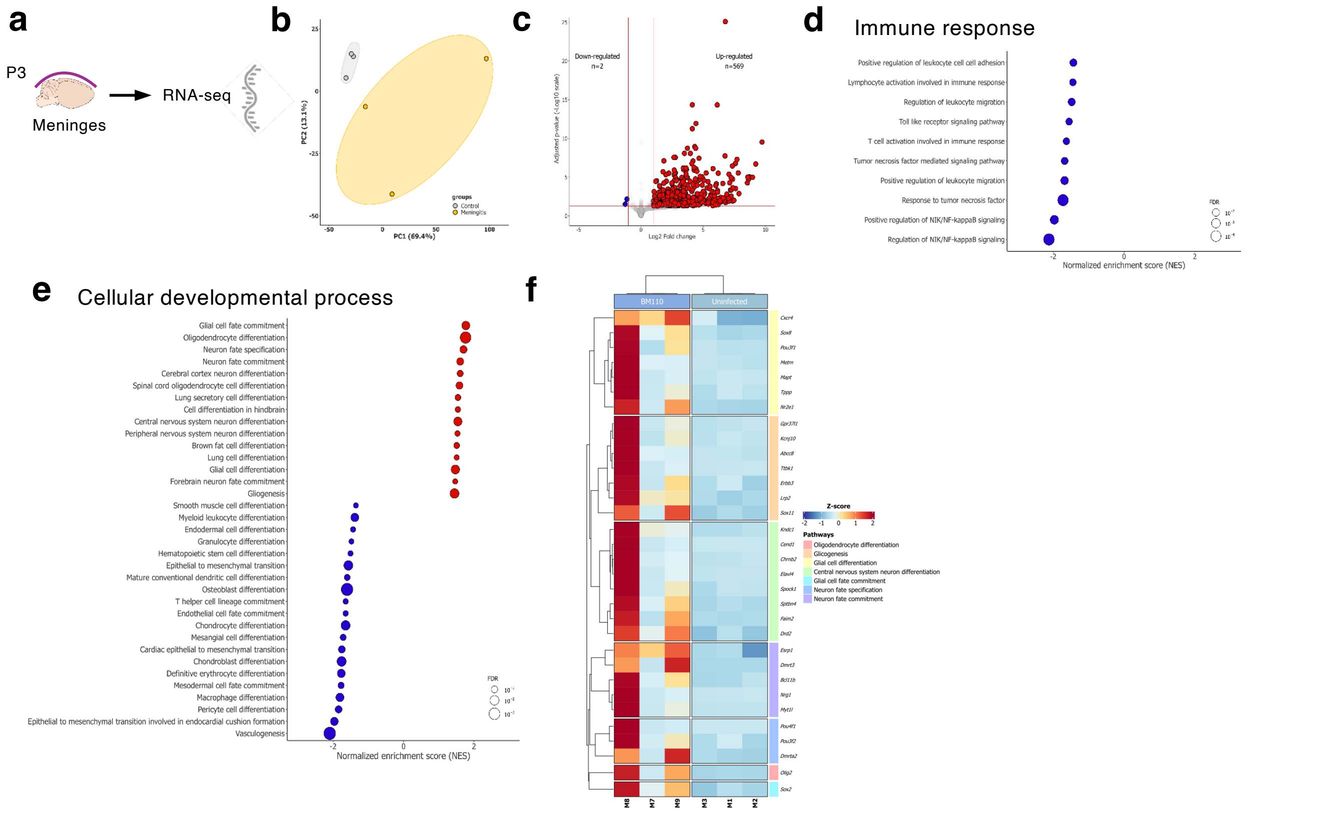

### Supplementary Figure 11

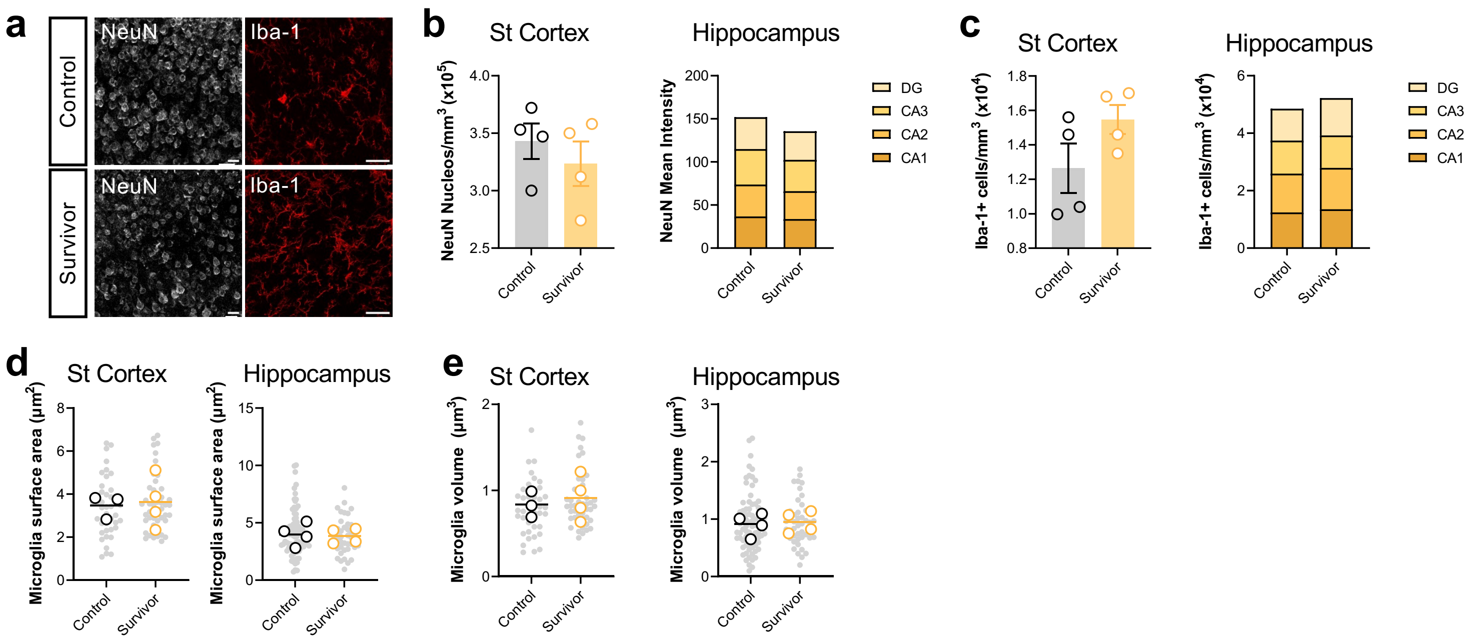
